# Global proteome metastability response in isogenic animals to missense mutations and polyglutamine expansions in aging

**DOI:** 10.1101/2022.09.28.509812

**Authors:** Xiaojing Sui, Miguel A. Prado, Joao A. Paulo, Steven P. Gygi, Daniel Finley, Richard I. Morimoto

## Abstract

The conformational stability of the proteome has tremendous implications for the health of the cell and its capacity to determine longevity or susceptibility to age-associated degenerative diseases. For humans, this question of proteome conformational stability has the additional complexity from non-synonymous mutations in thousands of protein coding genes challenging the capacity of the proteostasis network to properly fold, transport, assemble and degrade proteins. Here, we quantify the proteome-wide capacity to such challenges using the isogenic organism *Caenorhabditis elegans* by examining the dynamics of global proteome conformational stability in animals expressing different temperature-sensitive (ts) proteins or short polyglutamine (polyQ) expansions in the context of biological aging. Using limited proteolysis of native extracts together with tandem mass tag-based quantitative proteomics, we identify proteins that become metastable under these conditions and monitor the effects on proteome solubility and abundance. Expression of different mutant proteins in the same tissue identifies hundreds to a thousand proteins that become metastable affecting multiple compartments and processes in a cell autonomous and non-autonomous manner. Comparison of the network of metastable proteins, however, reveals only a small number of common proteins. The most dramatic effects on global proteome dynamics occur in aging with one-third of the proteome undergoing conformational changes in early adulthood. These age-dependent metastable proteins overlap substantially with ts proteins and polyQ; moreover, expression of polyQ accelerates the aging phenotype. Together, these results reveal that the proteome responds to misfolding one-at-a-time to generate a metastable sub-proteome network with features of a fingerprint for which aging is the dominant determinant of proteome metastability.

## INTRODUCTION

Biological systems are distinguished by their dynamics, robustness and stress resilience. For protein biology this is determined by the relationship between the physicochemical properties of proteins to fold stably into functional structures, the cell stress responses that detect and respond to the appearance of damaged proteins and the protein homeostasis (proteostasis) network that is essential in the cell to fold, refold or degrade the proteome^1–3^.

Within a typical cell, this involves many proteins assembled as macromolecular machines to transmit genetic information, generate energy, movement and response to the environment and organized into metabolic and sensory transducing networks. For metazoans, the richness of protein biology generates the cellular diversity for tissue and organismal function, cellular health and longevity. The challenge, however, is with the complexity from millions of parts in the face of sequence variation from 10-13,000 non-synonymous coding mutations as occurred among individuals within human population^4,5^. Such mutations can contribute to protein conformational diversity and affect folding, generate alternative conformations, novel protein-protein interactions and subcellular localization or affect protein half-life^6,7^. Consequently, coding polymorphisms are the basis for cellular dysfunction as occurred in age-associated protein conformational diseases^8^.

Such polymorphisms contribute directly to the genetic basis of disease as reflected by the vast number of dominant and recessive genetic diseases with complex phenotypes such as Huntington’s disease (HD) that has been defined by expanded polyglutamine (polyQ) repeats, yet polyQ only accounts for 40 – 60% of the variation in age of onset with the remaining attributable to genetic factors other than the Huntingtin gene or environmental factors^9–12^. At the other extreme are Alzheimer’s disease and related dementias for which explanations other than a single dominant gene are necessary^13–16^. HD, as for all other neurodegenerative diseases, is a proteinopathy associated with polyQ length dependent aggregation^17–25^. In human disease and in invertebrate and vertebrate animal model systems, aggregation and toxicity of mutant Huntingtin with expanded polyQ is age-associated^18,20,21,26–29^. Studies in *Caenorhabditis elegans* (*C. elegans*) have shown that expression of polyQ across the lower threshold of Q35-Q40 or mutations in superoxide dismutase 1 (SOD1) associated with amyotrophic lateral sclerosis (ALS) in neurons or muscle cells causes the misfolding and loss-of-function of co-expressed metastable folding sensors that exhibit *in vivo* temperature-sensitivity (ts)^30,31^. Furthermore, the polyQ-induced misfolding of expressed ts proteins at the permissive condition increases the aggregation load in the cell to enhance polyQ aggregation and proteotoxicity^30^. Unlike polyQ, mutant SOD1 proteins, though highly aggregation-prone in *C. elegans*, exhibits reduced intrinsic proteotoxicity^31^. The demonstration that toxicity of both mutant SOD1 and polyQ expansions can be modulated by metastable proteins supports the contention that the proteostasis network is sensitive to cumulative protein damage, and that the disruption of protein folding may be a common mechanism that underlies the toxicity of aggregation-prone proteins.

The use of ts proteins as folding sensors also revealed that aging alone was sufficient to promote misfolding and aggregation and established a formal link between the environment and protein folding thermodynamics at the permissive condition^32^. Proteostasis collapse in aging is not random but highly regulated by signals from the germline stem cells to repress activation of the heat shock response and other organellar cell stress responses at reproductive maturity leading to failure in quality control and an imbalanced proteome during somatic aging^33^. The accumulation of misfolded proteins, in turn, contributes to the progressive disruption of the folding environment when this balance becomes overwhelmed, e.g., by the expression of an aggregation-prone protein in conformational diseases^30^. Therefore, the presence of marginally stable or folding defective proteins in the genetic background of conformational diseases act as potent extrinsic factors that modify aggregation and toxicity^30,31^. Given the prevalence of polymorphisms in the human genome^5^, they could contribute to variability of disease onset and progression. This interpretation also provides a mechanistic basis to the notion that the late onset of protein misfolding diseases may be due to gradual accumulation of damaged proteins, resulting in a compromise in folding capacity.

This hypothesis offers an explanation for the apparent paradox of cell-type-specific toxicity caused by ubiquitously expressed toxic proteins in conformational diseases, such as SOD1-related ALS, HD, Alzheimer’s disease and related dementias. The differential modulation of mutant SOD1 toxicity in *C. elegans* by specific ts mutations and the interplay with expanded polyQ suggests that the presence of mildly destabilizing protein polymorphisms in the genetically diverse human population could direct such specific phenotypes^30,31^. The expression of a distinctive complement of expressed proteins in a particular tissue would generate cell-type specific genetic interactions among aggregation-prone proteins with destabilizing polymorphisms. In turn, this would be predicted to generate disease variability across the human population with collections of protein polymorphisms specific to individuals or families, and therefore missed in the population-based studies^34^. Up to 70% of rare missense alleles are mildly deleterious in humans^35^; some of these polymorphisms may result in the production of metastable or folding-deficient proteins. Identification of cell-specific pathways or protein complexes, which may dysfunction when folding or stability of their components is challenged by co-expression with an aggregation-prone protein, may thus provide specific toxic mechanisms for conformational diseases and help focus the search for the disease-modifying polymorphisms.

Here, we examine the dynamics of global proteome conformational stability in animals expressing different ts proteins or short polyQ expansions in the context of biological aging. We addressed this using limited proteolysis coupled with tandem mass tag-based quantitation by mass spectrometry (LiP-TMT-MS) to identify proteins that become conformationally metastable with effects on proteome solubility and abundance. Expression of different ts proteins in the same tissue causes hundreds of proteins to become conformationally metastable at the restrictive condition affecting multiple compartments and processes with cell non-autonomous effects. Comparison of these metastable proteins, however, reveals only a small number of common proteins and condition-specific expression of small heat shock proteins (sHSPs) and specific HSP70 chaperones distinct from the canonical heat shock response. Analysis of animals expressing polyQ24, Q35 and Q40 at the threshold of aggregation identifies a polyQ-dependent metastable sub-proteome of a distinct set of hundreds of proteins. Aging has the most dramatic effects on global proteome dynamics with one-third of the proteome undergoing conformational changes, mostly between days 1 to 6. These age-dependent conformationally metastable proteins overlap substantially with ts proteins and polyQ; moreover, expression of polyQ accelerates the aging phenotype.

## RESULTS

### Application of limited proteolysis coupled with tandem mass tag mass spectrometry (LiP-TMT-MS) to examine global protein conformational metastability at the organismal level

*C. elegans* as a biological animal model system has the distinctive feature of being isogenic, therefore allowing us to investigate global proteome metastability in response to the effects of a single coding polymorphism, the effects of expanded polyQ proteins and aging. Here, we have taken advantage of well-studied ts missense mutations to examine their proteome-wide effects on protein metastability in animals at both the restrictive (25°C) and permissive (15°C) conditions. As ts mutations are themselves metastable, we reasoned that they should have effects on protein conformation that can be detected using reported LiP-MS^36^. LiP-MS relies on the use of a non-specific protease, proteinase K (PK) to detect conformationally accessible regions of proteins under native conditions, followed by trypsin digestion on denatured fragments to produce shorter peptides that are compatible with traditional proteomics. This method combined with TMT isobaric labelling (LiP-TMT-MS) (Fig. 1A) allows an accurate quantitation and reproducible comparison for up to 18 samples per single MS experiment^37,38^. It also reduces the number of missing values across all samples otherwise observed in other MS methods.

**Figure 1.**
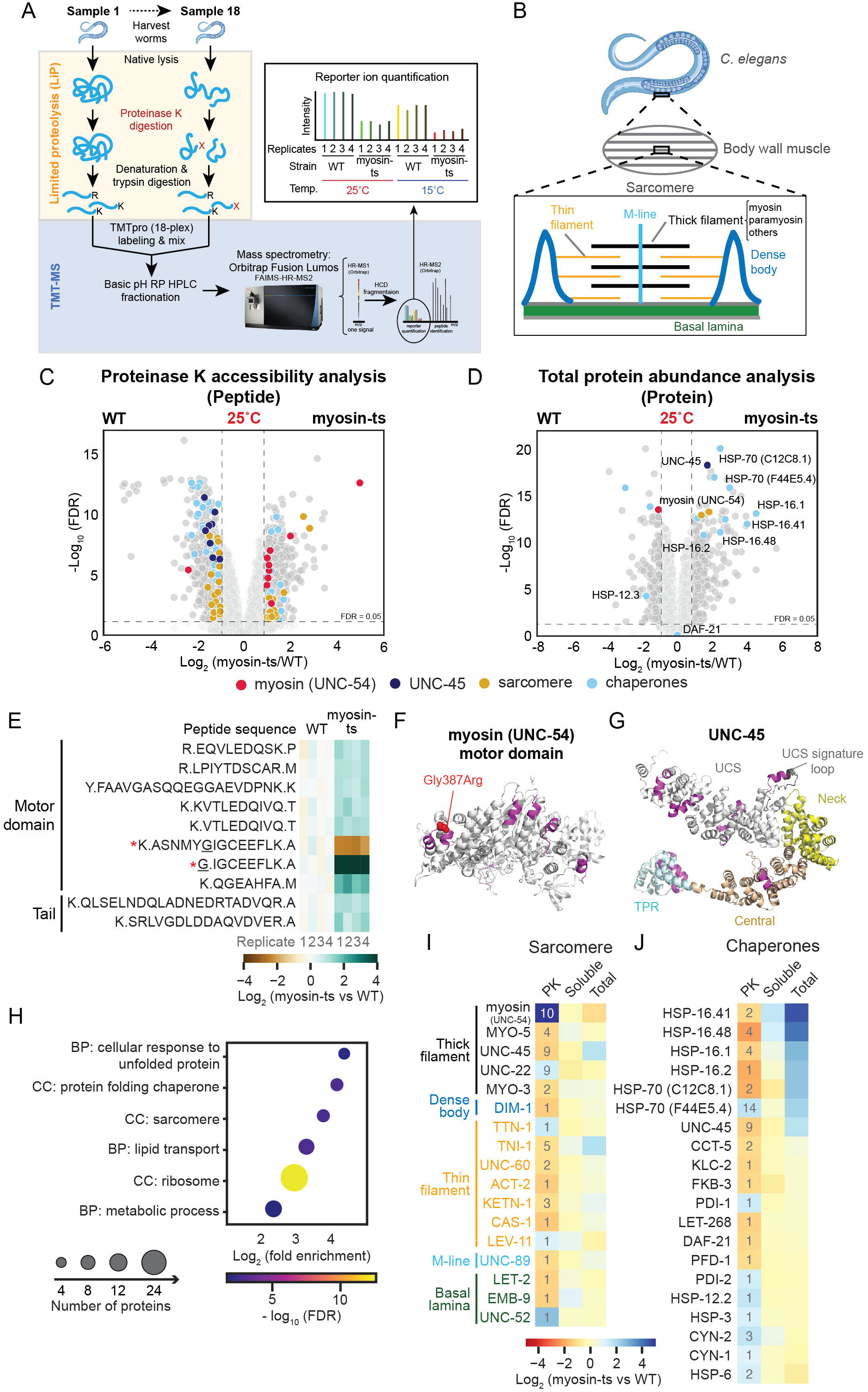
Impact of a single temperature-sensitive (ts) missense mutation in myosin (UNC-54) on the conformational metastability of endogenous proteins at the restrictive condition. (A) Proteomic workflow for limited proteolysis (LiP) coupled with tandem mass tag (TMT)-based quantitative measurements of protein conformational changes across different conditions (up to 18 samples). Native protein lysates were subjected to proteinase K (PK) digestion for 1 min. Proteins with conformations that are more accessible to PK (e.g., Sample 18) are more likely to be cut, therefore generating structure-specific fragments. After denaturation, these fragments were further digested by trypsin into smaller peptides. Fully-tryptic (shown as -K or -R) and half-tryptic (shown as -X, indicating amino acids that are not K, R or C-terminal) peptides were labelled with TMT and mixed. The mixtures were fractionated by basic pH reversed-phase high performance liquid chromatography (basic pH RP HPLC) to reduce the proteome complexity. The fractionated mixtures were subject to mass spectrometry (MS) analysis in an instrument equipped with a FAIMS pro interface. TMT reporter ions were detected in the orbitrap (HR-MS2) after HCD fragmentation. After database search, peptides and proteins were identified. The total peptide intensities of each TMT channel were normalized to be equal. Peptide ratios were calculated based on the normalized TMT reporter ion intensities. To control for protein abundance changes between conditions, the peptide ratios were corrected by their corresponding protein ratios generated from the trypsin-only (TO) pipeline from Fig. S1A. From here on, peptide ratios from PK accessibility analysis refer to the corrected ratios (for correction details, see the Materials and Methods session). (B) Cartoon of the organization of sarcomere in the body wall muscle cells of *C. elegans*. Sarcomere is constituted of thick filament (e.g., myosin, paramyosin, colored in black), thin filament (orange), M-line (light blue), dense body (dark blue), and basal lamina (green). (C – D) Volcano plots showing the comparisons of peptides from PK accessibility analysis (C) and proteins from total abundance analysis (D) between myosin-ts and WT animals grown at the restrictive condition (25°C) at day 1 of adulthood. Peptides (C) and proteins (D) with at least two-fold change and false discovery rate (FDR) < 0.05 (both thresholds indicated with dash lines) were considered as significant changes (colored in dark grey). The FDR was obtained from linear models for microarray (LIMMA) analysis using Benjamin-Hochberg correction. Among the significantly changed peptides (C) and proteins (D), those belonging to myosin (UNC-54), UNC-45, other sarcomere, and chaperones were colored in red, dark blue, orange, and light blue, respectively. Proteins (D) were selectively annotated with their names. (E) Heatmap showing the myosin peptides with significantly differential PK accessibility between myosin-ts and WT animals from four biological replicates. * Represents those peptides that harbors the ts mutation (G387R, underlined). Note: the mutation site is cut by trypsin, therefore interfering the PK accessibility analysis. (F) Crystal structure of the myosin motor domain (PDB: 6QDJ^84^) with all peptides with significant changes in PK accessibility colored in purple. The mutation site (G387R) is highlighted in red. (G) Crystal structure of UNC-45 (PDB: 6QDK^84^) with all peptides with significant changes in PK accessibility colored in purple. The UCS domain was colored in white with the signature loop in grey, central domain in wheat, neck in yellow, and TPR domain in cyan. (H) Dot plot showing the Gene Ontology (GO) enrichment of biological process (BP) and cellular component (CC) for proteins with significant changes in PK accessibility between myosin-ts and WT animals grown at the restrictive condition. X-axis indicates logarithm of fold enrichment of GO terms. Dot color indicates FDR values calculated from Fisher’s exact test using FDR correction for multiple testing. Dot size indicates the number of proteins from our dataset that are included in the enriched GO term. For full list of enriched GO terms, see Table S2. (I – K) Heatmaps showing the changes in PK accessibility, solubility, and total protein abundance of sarcomeric proteins (I) and chaperones (K) that showed significant changes in PK accessibility between myosin-ts and WT (from Fig. 1C). The number shown under the PK column indicates the number of peptides with significant changes in PK accessibility quantified for the protein indicated in each row. The median values of fold-changes of these peptides represents the coloring in the PK column. Protein names (I) were colored based on their localisation on the sarcomere (same as Fig. 1B, thick filament: black, thin filament: orange, M-line: light blue, dense body: dark blue, and basal lamina: green).

Native worm lysates were subjected to a limited concentration of PK for a brief treatment, and subsequently denatured and digested with trypsin. Regions of a protein that are conformationally exposed will be more accessible for PK digestion, resulting in abundance changes in fully- or half-tryptic peptides (Fig. 1A). To control for changes in protein abundance between conditions, the same protocol without PK digestion, is performed in parallel (Fig. S1A, left panel). This analysis, called trypsin-only (TO) digestion, was accomplished on the soluble lysates, therefore providing information on protein abundance changes in the soluble fraction. As a complimentary approach for protein solubility measurements, we also used a well-established ultracentrifugation-based cell fractionation method^39,40^ to quantify the protein level changes in the pellet fraction (Fig. S1A, right panel). In addition, to study total protein abundance changes, SDS-solubilized proteomes across multiple conditions were compared in parallel (Fig. S1A, middle panel). The peptides obtained from the four pipelines (PK, TO, aggregate and total proteome analysis) were TMT labelled, mixed into four independent sets, and analysed by MS (Figs. 1A and S1A). Relative peptide abundances were determined based on their corresponding TMT reporter ion intensities collected in the MS. We set the threshold of at least two-fold change and a false discovery rate (FDR) less than 0.05 for a significant change in PK accessibility, solubility, or total abundance, unless otherwise stated.

### A single ts missense mutation causes widespread protein metastability at the restrictive temperature

As a test of our hypothesis, we selected myosin heavy chain B, a major component of the body wall muscle cell thick filament (Fig. 1B), and the well-studied missense mutation G387R (*unc-54(e1301)*)^41^, referred to hereafter as myosin-ts. Myosin-ts animals exhibit an uncoordinated loss-of-motility phenotype at the restrictive condition and are fully functional at the permissive condition through day 1 of adulthood^30,32,41^. We compared the proteome changes in PK accessibility, solubility (TO) and total abundance between myosin-ts and WT in day 1 adult animals at the restrictive and permissive conditions using TMT-16 plex labelling with four biological replicates (Fig. 1A, Table S1). With high reproducibility between replicates (Fig. S1B), we quantified 8,062 proteins in total from the three separate TMT experiments, including 5,752 proteins and 83,586 peptides from LiP-TMT-MS experiment (Figs. 1C and D; for protein quantification numbers in TO and total proteome experiments, see Table S1). This yielded 1,086 peptides corresponding to 454 proteins that showed significant changes in PK accessibility in myosin-ts mutant animals compared with age-matched WT at the restrictive condition (Fig. 1C). As myosin G387R is misfolded at the elevated temperature, this was used to demonstrate the sensitivity of LiP-TMT-MS to detect protein conformational changes. At the restrictive condition, ten peptides in myosin-ts with significant changes in PK sensitivity were identified (Fig. 1E) of which eight peptides are within the myosin motor domain that contains the G387R ts mutation (Fig. 1F) with two additional peptides identified in the tail region. Associated with these conformational effects on myosin-ts protein is a two-fold reduction of myosin total levels (Fig. 1D).

As myosin specifically interacts with the myosin co-chaperone UNC-45 during its folding and assembly in the sarcomere^42^, we asked whether LiP-TMT-MS detects changes in these interactions. UNC-45 is a multidomain protein that contains the myosin-binding UNC-45/CRO1/She4 (UCS) domain, neck, central and tetratricopeptide repeat (TPR) domain^42^. At the restrictive condition in which myosin-ts is dysfunctional, we detected nine peptides of UNC-45 that exhibited differential PK accessibility of which five mapped to the UCS domain (Fig. 1G) that includes a peptide identified in the UCS signature loop critical for myosin binding^42^. These results reveal altered interactions between myosin-ts and UNC-45 consistent with the disruption of myosin-ts folding and function at the restrictive condition.

Gene Ontology (GO) analysis of the 454 proteins that exhibited myosin-ts dependent PK sensitivity changes revealed an enrichment for components of the sarcomere, protein folding chaperones, ribosome, lipid transport and metabolic processes (Fig. 1H, Table S2). Within the sarcomere, we identified 17 muscle proteins in myosin-ts at the restrictive condition including proteins in the thick filament (UNC-54, UNC-45, MYO-5, MYO-3, UNC-22), thin filament (TTN-1, TNI-1, UNC-60, ACT-2, KETN-1, CAS-1, LEV-11), M-line (UNC-89), dense body (DIM-1) and basal lamina (LET-2, EMB-9, UNC-52) with minimal effects on solubility or abundance (Fig. 1I). These results suggest that the misfolding of myosin itself leads to widespread conformational metastability throughout the sarcomere. Another functional group that is highly enriched for PK sensitivity are chaperones and co-chaperones^43^ including four sHSPs (i.e., HSP-16.41, HSP-16.48, HSP-16.1 and HSP-16.2), two stress-inducible HSP70s (C12C8.1 and F44E5.4), and the myosin-specific chaperone UNC-45. The abundance of these chaperones increased by 2 – 23-fold (Fig. 1D and 1J). It is notable that we only detected one peptide changing in PK sensitivity and no change in the abundance of HSP90 (DAF-21) which is essential for the folding of myosin (Fig. 1D and 1J).

### Extensive changes in protein conformation also occur in animals expressing a ts mutation at the permissive condition

A critical control to interpret the proteome metastability generated by myosin-ts at the restrictive condition is to determine if any changes in proteome metastability occur in the same animals at the permissive condition where no discernable phenotype is detected. Proteomics analysis of myosin-ts animals compared to WT animals at the permissive condition identified 162 proteins with significant changes in PK accessibility (Fig. 2A) and 53 proteins exhibited changes in total abundance (Fig. 2B). GO analysis of PK accessible proteins revealed an enrichment for protein folding chaperones, glycolytic processes, and lipid transport (Fig. 2C, Table S2). Examination of the sarcomeric proteins revealed that myosin did not change in protein abundance at the permissive condition, although one myosin peptide (excluding the peptides containing the ts mutation) had a 1.8-fold change in PK accessibility (Figs. S2A and B). This suggests that myosin-ts animals at the permissive condition are pseudo-WT despite there being no behavioral consequence on motility or morphological effects on myofilament structure^32,41^.

**Figure 2.**
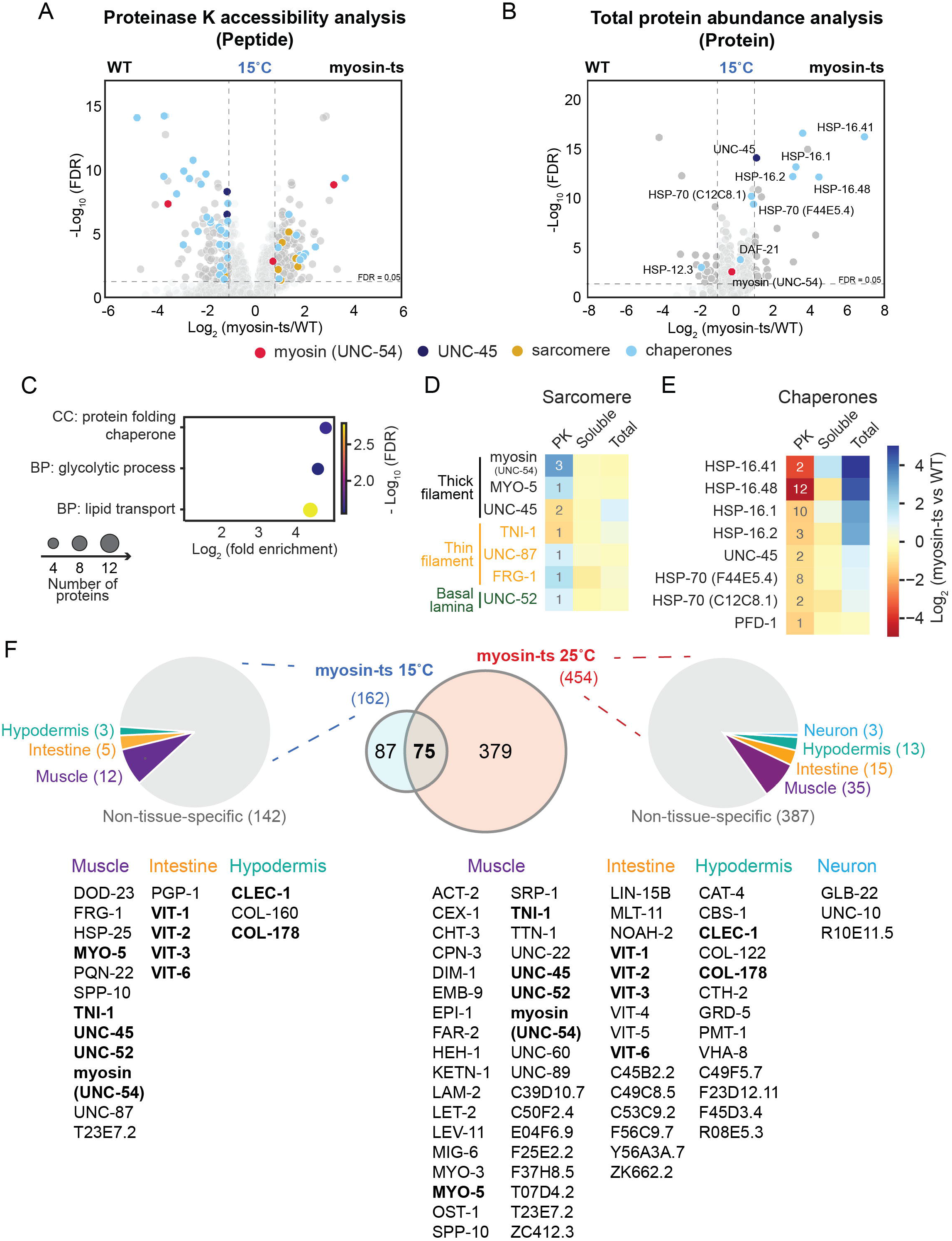
Impact of a single ts missense mutation in myosin on the conformational metastability of endogenous proteins at the permissive condition at day 1 of adulthood. (A – B) Volcano plots showing the comparisons of peptides from PK accessibility analysis (A) and proteins from total abundance analysis (B) between myosin-ts and WT animals grown at the permissive condition (15°C) at day 1 of adulthood. Peptides (A) and proteins (B) with at least two-fold change and FDR < 0.05 (both thresholds indicated with dash lines) were considered as significant changes (colored in dark grey). The FDR was obtained from LIMMA analysis using Benjamin-Hochberg correction. Among the significantly changed peptides (A) and proteins (B), those belonging to myosin (UNC-54), UNC-45, other sarcomere, and chaperones were colored in red, dark blue, orange, and light blue, respectively. Proteins (B) were selectively annotated with their names. (C) Dot plot showing the GO enrichment of BP and CC for proteins with significant changes in PK accessibility between myosin-ts and WT animals grown at the permissive condition. Dot color indicates FDR values calculated from Fisher’s exact test using FDR correction for multiple testing. Dot size indicates the number of proteins from the selected dataset that are included in the GO term. For full list of enriched GO terms, see Table S2. (D – E) Heatmaps showing the changes in PK accessibility, solubility, and total protein abundance of sarcomeric proteins (D) and chaperones (E) that showed significant changes in PK accessibility between myosin-ts and WT (from Fig. 2A). The number shown under the PK column indicates the number of peptides with significant changes in PK accessibility quantified for the protein indicated in each row. The median values of fold-changes of these peptides represents the coloring in the PK column Protein names (A) were colored based on their localisation on the sarcomere (same as Fig. 1B, thick filament: black, thin filament: orange, M-line: light blue, dense body: dark blue, and basal lamina: green). (F) Cell non-autonomous effects of misfolding of myosin-ts on metastable proteins in other tissues. Venn diagram shows the overlaps of proteins with significant changes in PK accessibility at the permissive and restrictive conditions. The pie charts show the tissue-specific mapping of the metastable proteins. The corresponding proteins in muscle, intestine, hypodermis, and neurons were listed alphabetically with those unassigned gene names at the bottom. Bold indicates proteins that are present both at the permissive and restrictive conditions. For full list of proteins, see Table S3.

These results further support the sensitivity of LiP-TMT-MS as a tool to detect protein conformational changes that either precede or are associated with phenotypic changes. Conformational changes were also observed in other sarcomeric proteins including the thick filament (MYO-5, UNC-45), thin filament (TNI-1, UNC-87, FRG-1) and basal lamina (UNC-52) consistent with myosin-ts at the permissive condition causing chronic molecular damage to the sarcomere (Fig. 2D). The class of proteins that exhibited the greatest change in PK accessibility and abundance are the four HSP16s, two HSP70s and UNC-45 (Figs. 2A, B and E) that showed an increased abundance of 1.8 – 121-fold. These chaperones increased their mRNA levels from the third-stage larvae (L3) and peaked at young adulthood and declined in early aging in myosin-ts compared with WT animals (Fig. S2C). Notably, no change was observed for HSP90 (DAF-21) which is essential for myosin folding. Together, these results reveal that even at the permissive condition that myosin-ts has substantial consequences on the global stability of the proteome and that the selective chaperone response is likely important to prevent myosin-ts from misfolding in development through early adulthood.

### Misfolding of myosin-ts has cell non-autonomous effects on proteome metastability

We have previously shown that expression of myosin-ts in muscle activates transcellular chaperone signaling, suggesting that the misfolding events in muscle are sensed in surrounding tissues^44^. Along this line, we asked whether myosin misfolding can alter the stability of proteins in other tissues. Of the proteins with changes in PK accessibility in myosin-ts animals at the restrictive condition, 66 of 454 proteins can be bioinformatically assigned to specific tissues^45^ of which 35 (52.2%) are muscle (including 17 sarcomeric proteins), 15 (22.3%) are intestinal (e.g., six vitellogenin proteins), 13 (19.4%) are hypodermal (e.g., collagen proteins COL-122 and COL-178) and 3 (4.4%) are neuronal (i.e., UNC-10, GLB-22 and R10E11.5) (Fig. 2F, Table S3). Likewise, at the permissive condition, 20 of 162 proteins can be assigned to specific tissues corresponding to 12 (60.0%) in muscle, 5 (25.0%) in intestine and 3 (15.0%) in the hypodermis. Many of these tissue-specific proteins (11/22) overlapped between the permissive and restrictive condition, of which five proteins (i.e., UNC-54, UNC-45, MYO-5, TNI-4 and UNC-52) were in muscle, four vitellogenin proteins (i.e., VIT-1/2/3/6) in intestine and two (i.e., CLEC-1 and COL-178) in hypodermis. These results reveal unexpectedly that myosin-ts has both cell autonomous effects in body wall muscle cells and cell non-autonomous effects on protein metastability in other tissues.

### Expression of a missense mutation in a different muscle protein affects the conformation of a distinct metastable subproteome

One explanation for the relatively large number of proteins that are conformationally metastable due to expression of myosin-ts is the possibility that the proteome encodes a common set of metastable proteins that readily misfold. This would be revealed, for example, by the analysis of another missense mutation in a different muscle protein expressed in the thick filament of the sarcomere. To address this, we examined paramyosin L799F (*unc-15(e1402)*)^46^, another well-studied ts mutation that exhibits the uncoordinated phenotype and myofilament disruption at the restrictive condition^32,46^. We quantified 7,949 proteins in total using the abovementioned four pipelines (i.e., PK, TO, aggregate and total proteome analysis), including 71,132 peptides corresponding to 5,347 proteins from LiP-TMT-MS analysis (for protein numbers in TO, aggregate and total proteome analysis, see Table S1).

At the restrictive condition, 2,082 peptides corresponding to 1,074 proteins (Fig. 3A) showed significantly differential PK accessibility between paramyosin-ts and WT animals on day 1 of adulthood. Altered total levels were observed for 494 proteins (Fig. S3A). We detected three peptides in paramyosin with significantly differential PK accessibility at the restrictive condition (Fig. 3A). Associated with changes in PK accessibility, mutant paramyosin levels increased (1.8-fold) in the aggregate fraction (Fig. S3C). At the permissive condition, 2,352 peptides with significant changes in PK accessibility corresponding to 1,157 proteins (Fig. 3B) were detected. A significantly lower number of proteins (159) exhibited changes in abundance (Fig. S3B). Two peptides in paramyosin showed changes in PK accessibility (Fig. 3B) with no change in solubility or total abundance (Figs. S3B and C).

**Figure 3.**
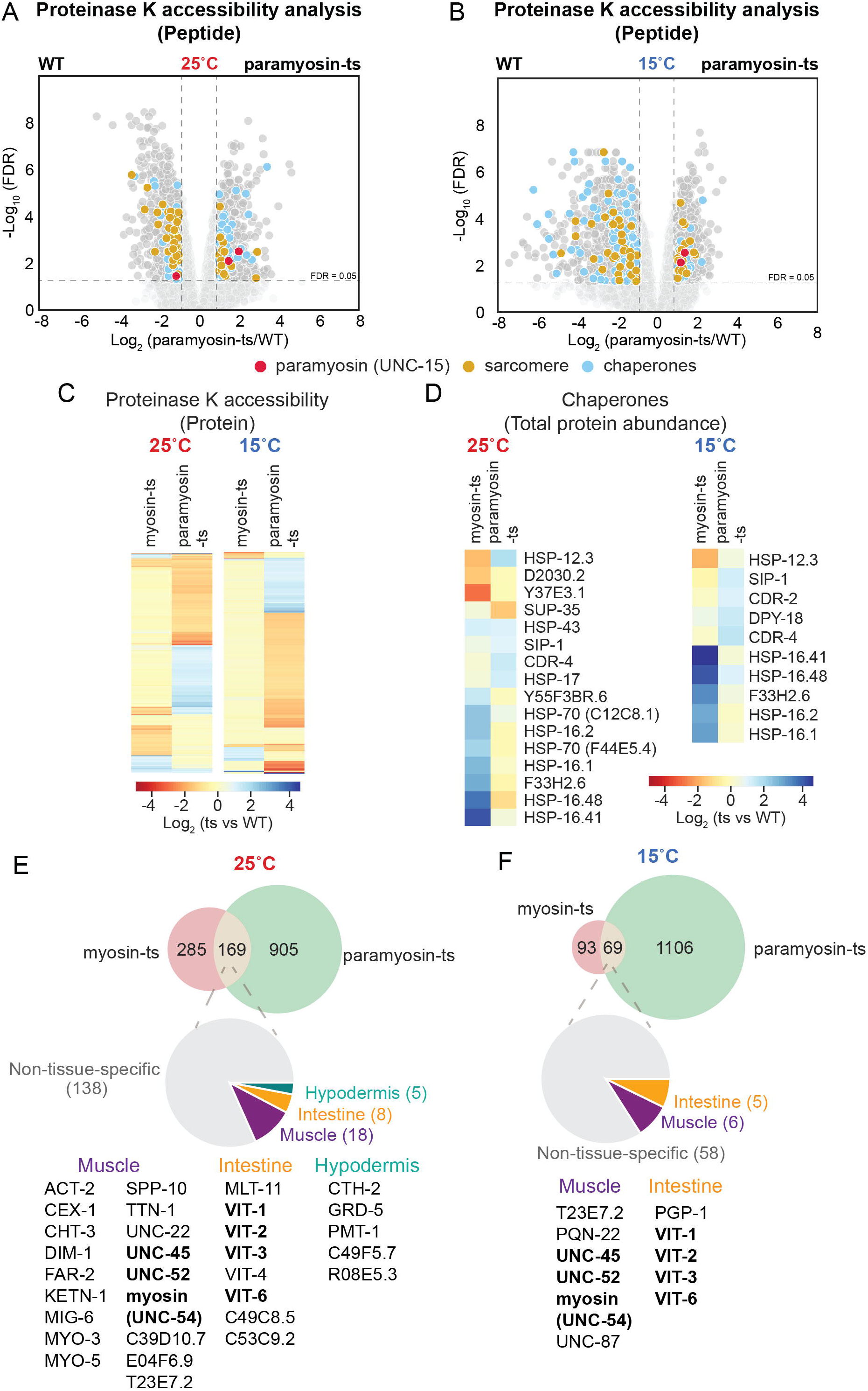
Distinct effects of paramyosin-ts on proteome conformational metastability compared with myosin-ts. (A – B) Volcano plots showing the comparison of peptides from PK accessibility analysis between paramyosin-ts and WT animals grown at the restrictive (A) and permissive (B) condition at day 1 of adulthood. Peptides with at least two-fold change and FDR < 0.05 (both thresholds indicated with dash lines) were considered significantly changed (colored in dark grey). The FDR was obtained from LIMMA analysis using Benjamin-Hochberg correction. Among the significantly changed peptides, those belonging to paramyosin (UNC-15), other sarcomere and chaperones were coloured in red, orange, and light blue, respectively. (C) Heatmap showing the distinct patterns of the changes of metastable proteins in myosin-ts and paramyosin-ts at the restrictive (left) and permissive (right) conditions. The median values of fold-changes of the peptides with significantly differential PK accessibility were represented for proteins in the graph. (D) Heatmap showing the upregulation at the total protein level of distinct chaperones in myosin-ts and paramyosin-ts at the restrictive (left) and permissive (right) conditions. (E and F) Cell non-autonomous effects of misfolding of myosin-ts and paramyosin-ts on metastable proteins in other tissues. Venn diagram showing the common metastable proteins between myosin-ts and paramyosin-ts at the restrictive (E) and permissive (F) conditions. The pie chart indicating the tissuespecific mapping of the common metastable proteins. The corresponding proteins in each tissue were listed alphabetically with unassigned gene names at the bottom. Protein names in bold indicates overlap with myosin-ts and paramyosin-ts analysis. For full list of tissue-specific proteins, see Table S3.

GO analysis of the metastable proteins induced by paramyosin-ts at the restrictive and permissive conditions showed enrichment for components in the sarcomere, lipid transport, proteasome complex, protein folding, translation and the lysosome (Fig. S3D, Table S4). Within the sarcomere, we identified PK accessibility changes in components of the thick filament (UNC-15, MLC-2, MLC-3, MYO-3, UNC-22, UNC-45 and myosin), thin filament (ACT-2, CAS-1, LEV-11, TNT-2, TTN-1, UNC-87), dense body (DIM-1), and basal lamina (UNC-52) at both conditions, with measurable effects in solubility and no major effects in total abundance (Figs. 3A, 3B and S3C).

Paramyosin-ts at the restrictive and permissive conditions had a nearly similar consequence on proteome metastability affecting 21.6% vs 20.1% respectively of the proteome. We then compared effects on proteome solubility. The metastable proteins were enriched in solubility changes at the restrictive condition (23.9%) compared with permissive condition (9.4%) (Fig. S3E), suggesting that the conformational changes at the restrictive condition are more likely associated with protein misfolding and aggregation.

### Comparison of the metastable sub-proteomes in animals expressing different missense mutations

Comparison of the conformationally metastable proteins in animals expressing paramyosin-ts with myosin-ts at both the restrictive and permissive conditions revealed distinct protein-specific patterns of changes (Fig. 3C). Paramyosin-ts has a much broader effect on proteome metastability than myosin-ts at both conditions (at 25°C, 1074 proteins vs 454 proteins; at 15°C, 1157 proteins vs 162 proteins, respectively). Analysis of the sarcomeric proteins showed a slightly higher proportion of proteins affected in paramyosin-ts (20) in comparison with myosin-ts (17) at the restrictive condition (Figs. 1I and S3C). This trend was more evident at the permissive condition with 20 proteins affected in paramyosin-ts and only 7 proteins affected in myosin-ts (Figs. 2E and S3C).

The distinct fingerprints of the metastable sub-proteomes between these two ts mutants were supported by the completely different profiles of chaperones upregulated in these two mutants (Figs. 3D). Myosin-ts dramatically upregulated major chaperones HSP70 and HSP16s, which are important to keep proteins folded and prevent protein aggregation. In contrast, only some sHSPs (i.e., HSP-43, HSP-17, SIP-1) were modestly upregulated in paramyosin-ts (Figs. 3D, S3A and B). The differential response of chaperones, in particular at the permissive condition, suggests that this is not a canonical heat shock response with coordinated upregulation of chaperones, but rather a specific and selective response to prevent further misfolding and potentially amplified proteotoxic damage at the permissive condition.

Examination of the tissue expression^45^ of common proteins affected by myosin- and paramyosin-ts provides further support for cell non-autonomous effects of misfolding of a muscle specific protein (Figs. 3E and F, Table S3). At the restrictive condition, 31 of 169 proteins (18.3 %) exhibited tissue-specific expression, of which 18 (58.0%) were muscle-specific with nine proteins localized to the sarcomere. Eight proteins (25.8%) including five vitellogenin proteins were intestine-specific and five were in hypodermis. Likewise, in animals at the permissive condition, 11 of 69 (15.9%) were tissue-specific; of which 63.6% overlapped with those at the restrictive condition, including sarcomeric proteins myosin, UNC-45 and UNC-52, and vitellogenin proteins VIT-1/2/3/6.

### PolyQ expansion proteins across the threshold of aggregation promotes proteome metastability

We had previously shown that expression of polyQ tagged with yellow fluorescent protein (polyQ-YFP) in body wall muscle cells led to misfolding at the permissive temperature of both myosin-ts (*unc-54(e1301)*) and paramyosin-ts (*unc-15(e1402)*) as a basis for polyQ proteotoxicity^30^. These experiments were limited by the ability to only monitor individual folding sensors, one-at-a-time, which did not allow a complete understanding of proteome-wide consequences of expression of expanded polyQ. To directly assess the effects of polyQ on the metastability of the entire proteome, we employed LiP-TMT-MS to identify the cohort of conformationally metastable proteins in *C. elegans* expressing Q24-YFP which is subthreshold for aggregation, and Q35-YFP and Q40-YFP that both exhibit age-dependent aggregation with different kinetics^29,47^. Moreover, because the polyQ-YFP transgenes are expressed under the control of the *unc-54* promoter in body wall muscle cells, this allowed the direct comparison to the metastable subproteomes in myosin-ts and paramyosin-ts animals.

LiP-TMT-MS analysis of subthreshold soluble Q24 compared to Q35 animals identified 258 peptides with altered PK accessibility corresponding to 128 proteins (Fig. 4A) while comparison of Q24 to aggregation-prone Q40 identified 1,744 peptides corresponding to 612 proteins (Fig. 4B). Of these, 80 of 128 (62.5%) metastable proteins in Q35 animals persisted in Q40 animals (Fig. 4C). This supports that there is a Q-length dependent accumulation of conformationally metastable proteins and are similar but not identical as the polyQ length expands. For example, Q35 affected the conformational stability of four proteins in the sarcomere (MLC-3, UNC-22, UNC-60 and PAT-10), which in Q40 animals expanded to include eight additional sarcomeric proteins including myosin, MYO-3, MLC-1/2 and TNI-3 (Figs. 4A and B), with no significant effects on their total abundances (Figs. S4A and B). This is further supported by GO analysis that showed an enrichment in the myosin complex (Fig. 4D, Table S5). Lipid transport was also affected in a Q-length dependent manner, with four proteins (i.e., VIT-4/5/6 and LPR-3) affected by Q35 and an additional six proteins (i.e., VIT-2, IPR-4/5, LEA-1, OBR-4 and ABT-2) in Q40 animals.

**Figure 4.**
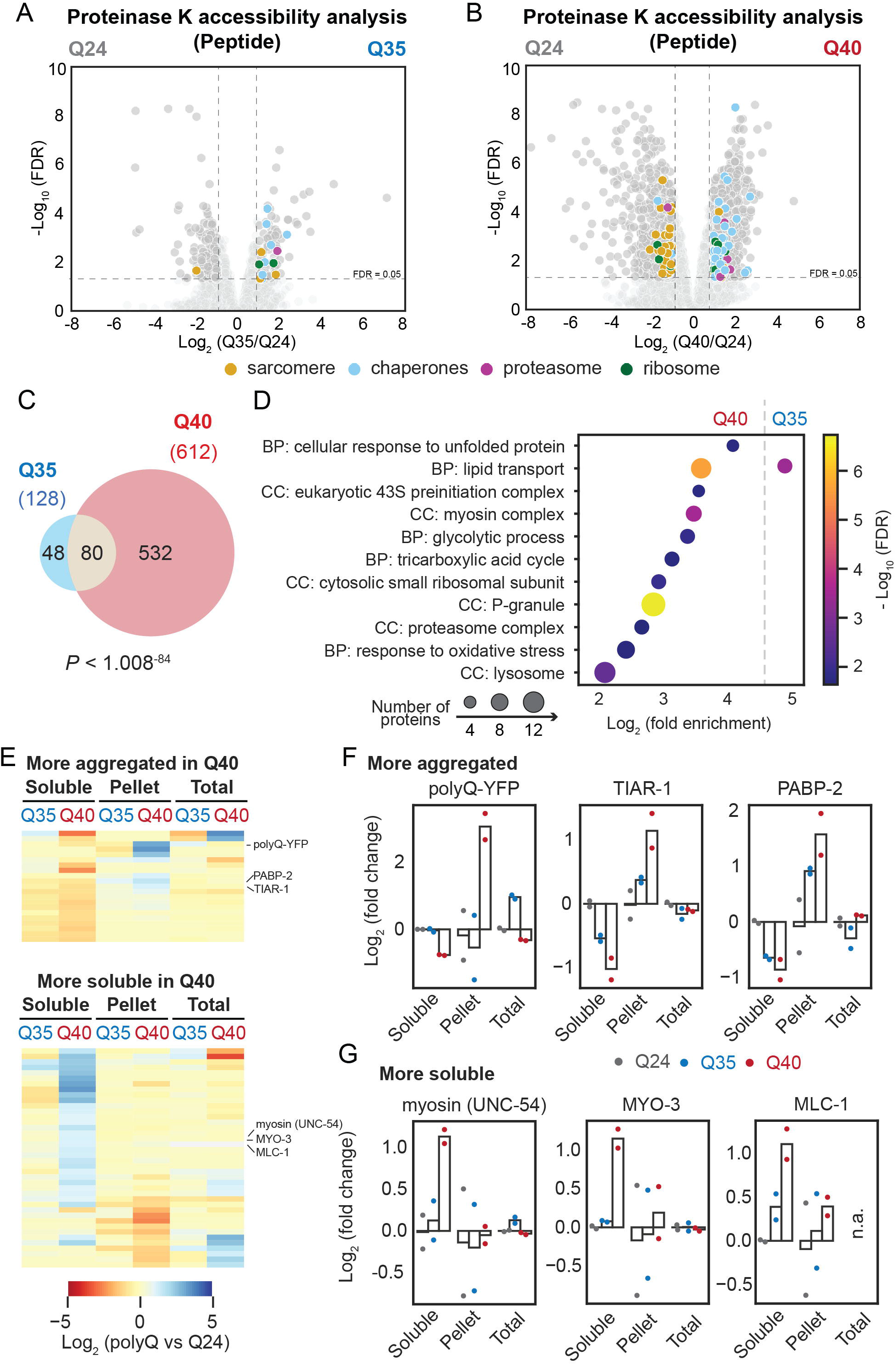
Q-length-dependent changes in protein conformational stability at day 1 of adulthood. (A – B) Volcano plots showing the comparison of peptides from PK accessibility analysis in Q35 vs Q24 (A) and Q40 vs Q24 (B) animals at day 1 of adulthood. Peptides with at least two-fold change and FDR < 0.05 (both thresholds indicated with dash lines) were considered significantly changed (colored in dark grey). The FDR was obtained from LIMMA analysis using Benjamin-Hochberg correction. Among the significantly changed peptides, those belonging to sarcomere, chaperones, proteasome, and ribosome were colored in orange, light blue, purple, and green, respectively. (C) Venn diagram showing the overlap of metastable proteins (with significant changes in PK accessibility) in Q35 and Q40, comparing with Q24 animals. The statistical significance was determined by Fisher’s exact test. (D) Dot plot showing the GO enrichment of BP and CC for proteins with significant changes in PK accessibility in Q35 and Q40 strains. Color indicates FDR values, and dot size indicates the number of proteins from the selected dataset that are included in the GO term. For full list of enriched GO terms, see Table S5. (E) Heatmap showing the metastable proteins that became more aggregated (upper) or more soluble (lower) in Q40, compared with Q24 animals. The protein abundance changes in soluble (TO), pellet and total fraction in Q35 and Q40 compared to Q24 were shown in the heatmap. Blue indicates increases in their abundances in the corresponding fraction, while orange indicates decreases in their abundances. Proteins not quantified were colored in grey. (F and G) Examples of metastable proteins from panel E that became more aggregated (F) or more soluble (G) in Q40 animals. Bar graphs represents the logarithm of fold change in Q24 (grey), Q35 (blue) and Q40 (red) in comparison to Q24. Log_2_ ratios of the soluble (TO), aggregate (pellet) and total protein abundance are represented. n.a. indicates not quantified in the corresponding experiment.

Of the proteins specifically affected by Q40, they were enriched in GO terms associated with protein quality control, including the proteasome, ribosomes, lysosomes, and stress responses including cellular response to unfolded protein and oxidative stress (Figs. 4A, B and D, Table S5). Other enriched GO terms included energy-related glycolytic process and tricarboxylic acid cycle, and P granules.

To address the functional implications of the conformationally metastable proteins in polyQ animals, we examined changes in solubility in Q35 and Q40 animals compared with Q24 control. In Q35 animals, only 3.1% (4 of 128) of metastable proteins had significant effects in solubility, which increased in Q40 animals to 10.0% (61 of 612). These include 21 proteins that increased in aggregation measured by the changes in the soluble and pellet fractions. Underscoring the robustness of the approach, polyQ, as well as core stress granule proteins that coaggregate with polyQ (i.e., TIAR-1, PABP-2)^48–50^ became more aggregated in Q40 animals (Figs. 4E and F). This is consistent with the findings that polyQ disrupts the dynamics of stress granules^40,51^. Forty proteins increased in solubility in Q40 animals including the myosin complex UNC-54, MYO-3 and MLC-1 (Figs. 4E and G), which suggests the disassembly of this macromolecular protein complex by Q40, consistent with the motility defect in Q40 animals^29^.

Next, we compared the metastable proteins in polyQ animals with genetic modifiers of polyQ (Q35 and Q37) aggregation previously identified through a genome-wide RNA interference screen in *C. elegans*^52^. There was no overlap between Q35 (Fig. S4C) and genetic modifiers and only 13 overlaps for Q40 (Fig. S4D). Despite the different timepoints (days 2 – 3 vs day 1 of adulthood) examined, the distinct set of metastable proteins may represent the diverse cascade events triggered by polyQ rather than the upstream regulatory pathways that modify polyQ aggregation. Comparing with the largest dataset of polyQ-expanded huntingtin exon 1 fragment (Httex1) interactors^51^ showed limited overlaps with metastable proteins in Q35 (20.3%) and Q40 (15.9%) (Figs. S4E and F, Table S6). The differences may be due to different biological systems and cell types (mouse Neuro2A cells vs *C. elegans*) used in the studies.

### Widespread increase in protein conformational metastability in early aging

Many studies have previously demonstrated extensive proteome-wide changes in *C. elegans* aging^39,53–56^. Building upon this, we measured the global effects of aging on protein conformation, in relation to changes in solubility and total abundance in days 1, 6 and 9 WT animals using the above-mentioned proteomic methods and pipelines (See Figs. S5A – C for comparisons of our total proteome analysis with previously published studies). We quantified 7,890 proteins from four TMT experiments (PK accessibility, TO, aggregate and total abundance), including 58,293 peptides corresponding to 5,252 proteins by LiP-TMT-MS (for protein numbers in TO, aggregate and total abundance analysis, see Table S1). Of these, 1326 proteins (25.2%) representing 3686 peptides exhibited significant changes in PK accessibility between day 1 to day 6 (Fig. 5A) that increased on day 9 by an additional 430 proteins (1199 peptides) corresponding to 33.3% of the detected proteome (Figs. 5B and C).

**Figure 5.**
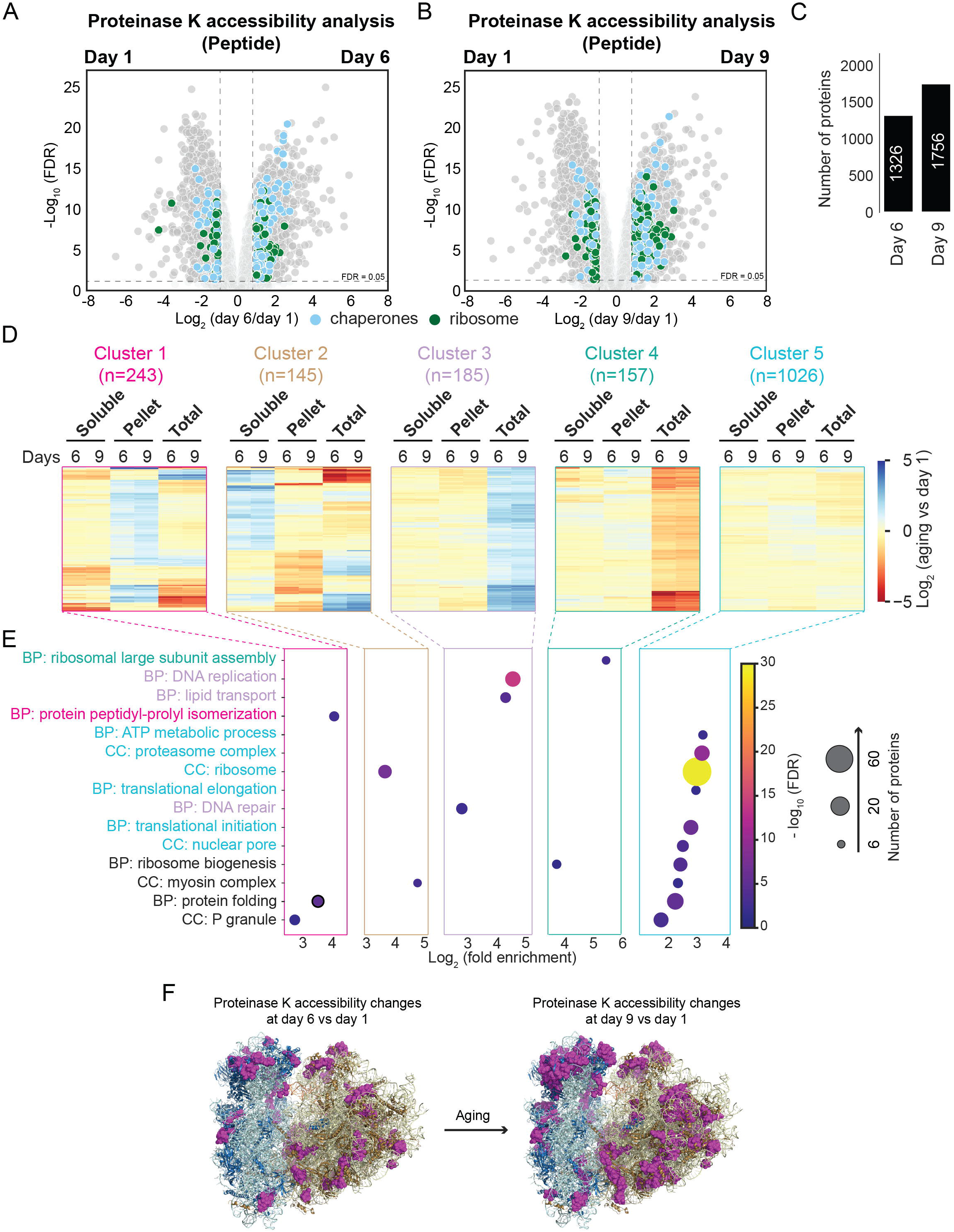
Widespread protein conformational changes in early aging in WT animals. (A – B) Volcano plots showing peptides with differential PK accessibility in day 6 vs day 1 (A) and day 9 vs day 1 (B). Peptides with at least two-fold change and FDR < 0.05 (both thresholds indicated with dash lines) were considered significantly changed (colored in dark grey). The FDR was obtained from LIMMA analysis using Benjamin-Hochberg correction. Among the significantly changed peptides, those belonging to chaperones were colored in light blue. (C) Bar plot comparing the number of proteins with differential PK accessibility between day 6 and day 9, both compared to day 1. (D) Heatmap illustrating the five clusters of conformationally metastable proteins according to their changes in solubility and total abundance between day 9 and day 1 in WT animals. Day 6 vs day 1 was also shown. Cluster 1, shown in pink, includes 243 proteins that became more aggregated (i.e., log_2_(fold change) > 1 in the pellet fraction or log_2_(fold change) < −1 in the soluble (TO) fraction, FDR < 0.05) in aging, regardless of their total abundance changes. Cluster 2, shown in brown, includes 145 proteins that became more soluble (i.e., log_2_(fold change) < −1 in the pellet fraction or log_2_(fold change) > 1 in the soluble fraction (TO), FDR < 0.05) in aging, regardless of their total abundance changes. Cluster 3, shown in light purple, includes 185 proteins that increased their total abundance (i.e., log_2_(fold change) > 1 in the total lysates, FDR < 0.05) without significant changes in their solubility (TO or pellet). Cluster 4, shown in green, includes 157 proteins that decreased their total abundance (i.e., log_2_(fold change) < −1 in the total lysates, FDR < 0.05) without significant changes in their solubility (TO or pellet). Cluster 5, shown in light green, includes 1026 proteins that only showed significant changes in PK accessibility in aging. The abundance changes in soluble (TO), pellet and total fractions were shown in the heatmap. (E) Dot plot showing the GO enrichment of BP and CC for proteins in five different clusters shown in (D). Cluster-specific GO terms were colored following the color scheme in (D), and shared GO terms were colored in black. Dot size indicates the number of proteins included in the GO term and dot color indicates FDR values. For full list of enriched GO terms, see Table S7. (F) Age-dependent changes in PK accessibility of ribosomal subunits. Cryo-EM structure of the human 80S ribosome (PDB: 6QZP^62^) with 40S subunits colored in blue, 18S rRNA in light blue, 60S subunits in wheat, 5S rRNA, 5.8 rRNA and 28S rRNA in light yellow, and tRNA in orange. All peptides with significant changes in day 6 vs day 1 (left panel), day 9 vs day 1 (right panel) in PK accessibility were colored in magenta.

To further understand the functional consequences of these age-dependent metastable proteins, they were sorted into five clusters according to their solubility and total protein levels profiles (Fig. 5D).

Cluster 1 (243 proteins) became more aggregated in aging regardless of their total abundance changes. They were enriched in P granules, protein folding, and protein peptidyl-prolyl isomerization (Fig. 5E, Table S7). Several P granule proteins (e.g., PGL-1, MEX-3) have been shown to form age-dependent aggregates^57^. Here we identified more uncharacterized RNA helicases (i.e., Y94H6A.5 and Y54G11A.3) that may be involved in reproductive aging. Protein peptidyl-prolyl isomerization is a rate-limiting step of protein folding and refolding^58,59^. The presence of peptidyl-prolyl cis-trans isomerases (i.e., PINN-1, CYN-2, FKB-4 and FKB-6) in the aggregates may lead to impairment of this process, which might be the basis of extensive protein conformational changes observed here.

Cluster 2 (145 proteins) became more soluble in aging, regardless of their protein abundance changes and were enriched in the myosin complex and ribosomes, consistent with dysfunction of these macromolecular complexes in aging^32,39,53,60,61^. We return to ribosomal subunits later.

Cluster 3 (185 proteins) increased in abundance without changes in solubility and were enriched in DNA replication, DNA repair and lipid transport. The metastability of DNA replication and repair could enhance the expression of damaged proteins that could further impair proteostasis. It’s notable that two sHSPs (i.e., HSP-16.1 and HSP-16.48) were also identified in this cluster (Figs. S5A and B), consistent with the age-dependent upregulation of these chaperones reported previously^39^.

Cluster 4 (157 proteins) decreased in abundance and were enriched in ribosome biogenesis and ribosomal large subunit assembly, consistent with the dysregulated ribosome biogenesis in aging previously observed^39,56^.

Cluster 5 (1,026 proteins) only exhibited changes in PK sensitivity without changes in solubility or abundance. These were enriched in a range of biological processes and cellular components including energy production (e.g., ATP metabolic process, glycolysis, fatty acid beta oxidation), protein quality control (e.g., protein folding, ribosome, proteasome complex, lysosome), translation (e.g., translational initiation and translational elongation), and nucleocytoplasmic transport (nuclear pore complex).

As ribosome-related processes were enriched in multiple clusters, we next pooled all ribosomal subunits for analysis. Mapping peptides with changes in PK accessibility to the ribosome structure^62^ revealed age-dependent metastability of ribosomal small and large subunits, clustered in 40S head and 60S (Fig. 5F). It’s notable that 59% (33 out of 56) of these metastable subunits became more soluble (1.5-fold – 5-fold) associated with decreases in their total abundances, including those localized in the 40S and 60S interfaces (Figs. S5D and E). Together, our data support a model that these subunits are dissociated from or unable to be assembled into the 80S ribosome complex, therefore causing loss of stoichiometry as a basis for ribosome dysfunction during aging^39^.

As aging affects translation demonstrated by our work and numerous studies^39,53,63,64^, we next asked whether the age-dependent metastable proteins represent those highly translated in aging, therefore vulnerable for misfolding. Cross comparison with age-dependent translatome in *C. elegans*^53^ showed that proteins with changes in PK accessibility had higher translation levels than those without at day 6 (Fig. S5F). Furthermore, a strikingly high fraction (91.8%) of the metastable proteins exhibited differential ubiquitination in aging^54^ while only 37.7% of them showed changes in phosphorylation, after filtering for commonly identified proteins in the datasets (Fig. S5F). Together, our data suggested that these metastable proteins are susceptible for damage, therefore imposing heavy burden on protein quality control machineries and impairing proteostasis in early aging.

### Is there a core set of conformationally metastable proteins across different protein misfolding stresses?

Here, we compared all proteins that exhibited differential PK accessibility under the conditions of animals expressing myosin-ts and paramyosin-ts at restrictive and permissive conditions, Q24, Q35, Q40, and aging on days 1, 6 and 9. In total, we identified changes in 3,008 proteins that exhibited significantly changed PK accessibility that corresponds to two-fifths of the proteome (39.1%). Despite this apparent widespread change in protein conformation, the proteins identified at risk were largely specific to the condition tested. Only four proteins (the vitellogenin proteins VIT-1, VIT-3, VIT-6 and a spliceosome structural protein SNR-2) exhibited altered protein conformation across all seven conditions (Fig. 6A, Table S8). Downregulation of VIT-1 has been implicated in enhanced lifespan^65^.

**Figure 6.**
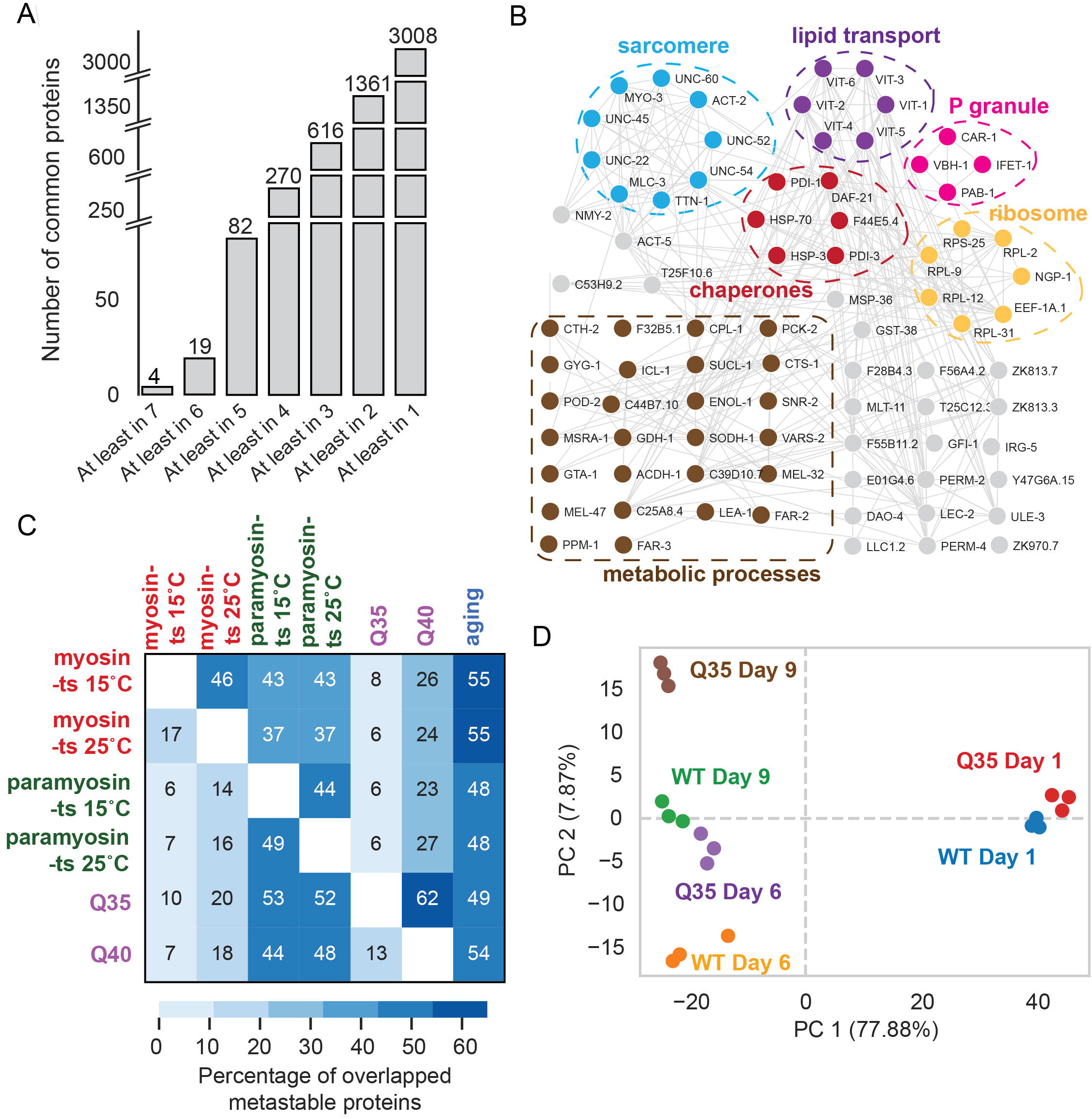
Comparison of conformationally metastable proteins in different mutants and aging. (A) Bar plot showing the number of proteins with changes in PK accessibility that were found in common by comparing different conditions. For full list of proteins, see Table S7. (B) Protein networks represented by STRING (v.11.0) analysis of proteins common in at least 5 conditions with medium confidence interactions. Selected significantly enriched GO terms are annotated. (C) Heatmap showing the percentages of overlapped proteins between comparisons. The percentage was calculated by the number of the overlapped proteins divided by the number of the metastable proteins in the mutant indicated on the left side of the heatmap. (D) Three biological replicates of Q35 and WT animals grown to days 1, 6 and 9 of adulthood were collected and analyzed for their changes in PK accessibility. PCA showed that different replicates of the same group cluster closely together in the two-dimensional plot (PC1 vs PC2), demonstrating the reproducibility among biological replicates. More importantly, this analysis also showed animals expressing Q35 at day 6 were clustered with WT animals at day 9.

As some proteins were not identified in all experiments due to different MS runs, we expanded the analysis to any six conditions. This resulted in 19 common proteins (including muscle-specific proteins myosin, FAR-2, intestinal proteins VIT-2, VIT-4, chaperone HSP70 (F44E5.4), enzymes involved in pyruvate metabolism (i.e., ICL-1, POD-2), glutamate dehydrogenase GDH-1, nucleolar GTP-binding protein 2 (NGP-1), and several uncharacterized proteins (ZK813.3, F28B4.3, T25C12.3, etc.) (Table S8).

Furthermore, in any five conditions, we observed 82 proteins (1% of the detected proteome, Table S8) with significant changes in PK accessibility. One of the enriched functional groups (Fig. 6B) is sarcomere, consistent with the perturbations initiated in the muscle. This also identified a core set of chaperones (i.e., HSP70 (C12C8.4 and F44E5.4), HSP90 (DAF-21), HSP-3, PDI-1 and PDI-3), suggesting a potential role for these chaperones to respond to a metastable proteome. Other major enriched GO terms included lipid transport and P granule, which further supports the cell non-autonomous effects of muscle protein misfolding on proteins in other tissues. We also detected changes in metabolic processes.

### Expanded polyQ accelerates age-dependent changes in proteome metastability

Comparing the effect of any two mutant proteins or conditions, we detected an overlap of 6% to 62% common metastable proteins (Fig. 6C). However, by cross-comparison with aging, consistently high overlaps (in the range of 48% to 55%) were observed, regardless of the mutations or conditions. This suggests that ts mutant animals at day 1 of adulthood or expression of polyQ promotes age-like proteome metastability.

To address this, we selected the chronic aggregation model of Q35 and examined its proteome metastability from day 1 to day 6 until day 9. The metastable proteome in day 6 Q35 animals formed a close cluster by principal component analysis with day 9 of WT animals (Fig. 6D), indicating the Q35 accelerates the age-dependent proteome metastability. Examining the total proteome abundance changes revealed a similar result (Fig. S6A). In agreement with this, increased levels of HSP-16.1 and HSP-16.48 were observed in Q35 animals at both day 6 and day 9, compared with age-matched WT animals (Fig. S6B). To understand the basis for the acceleration of the proteome clock by Q35, we performed the functional analysis of metastable proteins specifically affected by Q35 from day 1 to 9 when compared to WT. GO enrichment analysis (Fig. S6C) revealed age-dependent and chronic disruption of biological processes by Q35. At days 1 and 6, the effects were mainly on lipid transport and muscle contraction, as expected as Q35 is expressed in muscle. At day 9, we observed profound effects on protein quality control processes, including protein folding, translation and proteasome complex. Other GO terms included tricarboxylic acid cycle and P granule. Together, these results indicate that Q35 aberrantly engages with protein quality control machineries, therefore reducing proteostasis capacity and leading to the accelerated protein metastability in aging.

## DISCUSSION

This study addresses fundamental biological questions on the conformational stability of the proteome, and reveals that when a single protein misfolds, the conformation of hundreds of other proteins in multiple tissues are affected with consequences on protein solubility, abundance. These effects on global proteome metastability are strongly influenced by temperature as a potent environmental modifier and aging. The use of *C. elegans* as the biological animal model system is a central feature of this analysis in which isogenic animals provide the genetic background to test the effects of individual sequence polymorphisms by expression of ts mutant proteins or short polyQ expansion proteins across the threshold for protein aggregation in aging. These features combined with the use of LiP-TMT-MS proteomics provides precision measures of conformational changes in the global proteome with outstanding technical reproducibility. From our results, we propose that the expression of an intrinsically metastable protein generates a complex metastable subproteome fingerprint with features of a conformational network rather than the proposition that there is a common set of proteins that are at risk for misfolding. Only upon introducing the effects of aging does it become clear that aging is the dominant modifier of proteome conformational stability that overlaps substantially with the deleterious consequences of individual polymorphisms.

In the design of these experiments, we purposefully selected ts proteins that are expressed only in muscle cells of *C. elegans*. Expression of either myosin-ts or paramyosin-ts, both essential components of the muscle sarcomere, indeed, disrupts muscle function at the restrictive temperature generating the uncoordinated phenotype which itself does not cause lethality. At both the restrictive and permissive conditions, the misfolding of myosin-ts or paramyosin-ts resulted in substantial collateral damage throughout the sarcomere as determined by the number of components affected. While the cell autonomous effect on misfolding within muscle cells could be expected, the effects on the metastability of proteins expressed in other tissues is unexpected. One implication of these observations is that misfolding of a protein expressed solely in one tissue has cell non-autonomous effects in other tissues, in line with an organismal level control of proteostasis. This form of transcellular signaling had been previously observed in that the misfolding of myosin-ts in muscle cells could be corrected either by overexpression of HSP90, the myosin essential chaperone either in muscle cells or in neurons and intestine^44^.

The comparison of the proteome response for each ts protein at the restrictive and permissive condition also revealed that each condition renders a chaperone-specific response distinct from a canonical heat shock response. This was most strikingly observed with myosin-ts at the permissive condition with the over 100-fold increase in the levels of sHSPs together with a highly selective increase at much lower levels of two HSP70s. sHSPs are regulated by the transcription factors DAF-16 and HSF-1, which are both required for the potentiation of longevity by the insulin signaling pathway DAF-2^66,67^. Yet, despite the 100-fold induction of sHSPs, the proteome in myosin-ts animals at the permissive condition is not the same as WT animals, which suggests a tolerance for protein misfolding. Worth noting is that the chaperone response to paramyosin-ts or polyQ are dissimilar and in neither case was induction of the major heat shock protein HSP90 observed, which is essential for myofilament function.

Our results on proteome dynamics in animals expressing ts proteins at the permissive condition are most revealing and suggests that there is a level of protein conformational noise that is tolerated with no apparent behavioral and physiological consequence. The absence of phenotype in animals with ts mutations at the permissive condition is therefore a pseudo-WT state that reflects the compensatory changes or tolerance of conformational noise. That each proteotoxic challenge resulted in a unique conformational fingerprint suggests a network relationship in which folding stability is highly dependent on the environment (temperature) and age.

How might the observations presented here using *C. elegans* be relevant to human conformational diseases of aging? The magnitude of proteome metastability observed from one single coding polymorphism indicates a constantly remodeled human proteome in the face of genetic variation and aging. We posit that the complex phenotypic variation in HD, AD and related dementia, Parkinson’s disease, ALS, and other neurodegenerative diseases is the sum of proteostasis collapse in aging and the tremendous genetic variations in expressed coding polymorphisms.

## MATERIALS and METHODS

### C. elegans strains and maintenance

*C. elegans* were grown on standard Nematode Growth Medium (NGM) plates seeded with *Escherichia coli* (*E. coli*) OP50 bacteria at 20 °C unless otherwise stated. The Bristol strain N2 was used as wild-type (WT). The ts mutants used were myosin-ts – CB1301[*unc-54(e1301)*] and paramyosin-ts – CB1402[*unc-15(e1402)*], obtained from the *Caenorhabditis* Genetics Center (CGC) (University of Minnesota). The polyQ expressed in muscle strains used were: Q24m – AM138[rmIs130[*unc-54p::q24::yfp*], Q35m – AM140[rmIs132[*unc-54p::q35::yfp*] and Q40m – AM141[rmIs133[*unc-54p::q40::yfp*].

### Synchronization of large populations of C. elegans

To obtain large populations of synchronized worms for proteomics, bleaching of gravid adults was performed. Briefly, worms were washed with M9 buffer (22 mM KH_2_PO_4_, 42 mM Na_2_HPO_4_, 86 mM NaCl). After centrifugation at 1100 rpm for 1 min at room temperature, roughly 50 μL worm pellet was resuspended in alkaline hypochlorite solution (2 mL of 5.65 – 6% NaOCl (Fisher, cat# SS290-1), 2.5 mL 1 M NaOH, 5.5 mL dH_2_O) for 4.5 min to dissolve adults and obtain eggs. The mixture was then quickly centrifuged at 1100 rpm for 30 s to collect eggs, followed by three washes in M9 buffer. Eggs were then maintained in M9 buffer without food for 16 hours for the strains of N2, AM138 and AM140 grown at 20°C, for 24 hours for N2 and CB1301 grown at 15°C and AM141(20°C), or for 48 hours for CB1402 (15°C) to allow hatching and prevent further development beyond the L1 stage. For strains of CB1301, CB1402 and AM141, their eggs were not all hatched. To separate L1s from eggs, the mixture of eggs and L1 larvae were first placed on a fresh 10-cm NGM plate seeded with OP50, followed by washing L1s away from eggs on the plates with M9 buffer. The concentration of L1s was determined by counting under the microscope. 4000 L1s were plated for each 10-cm plate. A total of 12,000 L1s were regarded as one biological replicate for each strain.

### Growth conditions

For myosin-ts, paramyosin-ts and their WT control N2 worms, L1 larvae were either grown at the permissive (15°C) or restrictive (25°C) conditions until day 1 of adulthood. For polyQ (Q24, Q35 and Q40) strains, L1 larvae were grown at 20°C until day 1 of adulthood. For the aging experiments of Q35 and WT worms, L1 larvae were grown at 20°C to days 1, 6 and 9. To avoid starvation, worms were re-plated to fresh OP50-seeded 10-cm plates every day after day 1 of adulthood, and at least ten rounds of wash and sedimentation with M9 buffer was used to remove larvae contamination. All worm samples were collected by washing in Buffer 1 (20 mM HEPES, 150 mM NaCl, 10 mM MgCl_2_) for at least 6 times to remove bacteria and larvae contamination. Roughly 150 μL worm pellet was placed in a 2 mL Eppendorf ‘safe-Lok’ tube pre-chilled in liquid nitrogen (LN) and snap frozen and kept at −80°C until future use.

### Sample preparation for limited proteolysis (LiP)

Frozen worm pellets were homogenized with 4 × 90 s × 30 Hz pulses in a Retsch MM400 mixer mill, chilling in LN between pulses. 600 μL ice-cold Buffer 1 was added to the lysates. The protein concentration was determined by bicinchoninic acid (BCA) assay (Thermo) and was adjusted to 3 μg/μL with ice-cold Buffer 1. The lysates were then divided into three fractions for LiP (300 μL), protein solubility (200 μL) (see the session of Cell fractionation by ultracentrifugation) and total protein abundance (100 μL, see the session of Cell fractionation by ultracentrifugation) measurements. The soluble lysates for LiP were obtained from clarification at 16,000 g for 10 min at 4°C. 250 μL of the supernatant was carefully transferred to a new icecold 1.5 mL Eppendorf tube without disturbing the pellet. The protein concentration of the soluble fraction was determined by BCA assay and adjusted to 1 μg/μL with ice-cold Buffer 1. 200 μg of the soluble protein lysates were designated as PK-treated samples and an equal amount of the protein lysates were regarded as trypsin-only (TO) samples as control for PK samples. Briefly, PK and TO samples were incubated at 25°C for 5 min. Immediately, PK from *Tritirachium album* (Sigma Aldrich) was added to the PK samples at an enzyme:protein mass ratio of 1:30 and incubated for exactly 1 min. No PK enzyme was added to the TO samples. The digestion was quenched by heating PK and TO samples in boiling water for 5 min in a water bath, followed by addition of sodium deoxycholate (DOC) (Sigma Aldrich) to a final concentration of 5% to the cooled down samples. Both PK and TO samples were subjected to reduction of cysteine residues by 12 mM 1,4-dithiothreitol (DTT) for 45 min at 37°C, followed by alkylation by 55 mM iodoacetamide for 45 min at 37°C in dark. All samples were then digested by lysyl endopeptidase LysC (Wako Chemicals) at an enzyme:protein mass ratio of 1:100 for 2 hours at 37°C at 600 rpm in a thermomixer. The mixtures were then diluted to 1% DOC with 100 mM triethylammonium bicarbonate (TEAB), followed by digestion with trypsin (Thermo) at the enzyme:protein mass ratio of 1:100 overnight. The digestion was quenched by addition of trifluoroacetic acid (TFA) to a final concentration of 2%.

### Cell fractionation by ultracentrifugation

The separation of insoluble fraction was performed as previously described^39,40^. Briefly, 200 μL worm lysates were resuspended in an equal volume of Buffer 2 (Buffer 1 + 8 mM EDTA, 2% (v/v) IGEPAL CA-630, 1:500 EDTA-free protease inhibitor tablet (Roche)). The lysates were placed in a pre-chilled ultracentrifugation tube and centrifuged at 500,000 g for 10 min at 4°C. 350 μL of the resultant supernatant was reserved as the supernatant sample. The pellet was washed three times with 500 μL Buffer 3 (Buffer 1, 4 mM EDTA, 1% (v/v) IGEPAL CA-630, 1:1,000 EDTA-free protease inhibitor tablet (Roche)). The supernatant and pellet fractions were then denatured by SDS to a final concentration of 2% SDS and 4 mM DTT, followed by incubation at 95°C for 20 min. The denatured protein samples were reduced and alkylated as described above. Protein lysates were then purified by methanol-chloroform precipitation^68^. The resultant pellet was resuspended in 100 μL 8 M urea, 50 mM TEAB with vigorous vortexing. The protein concentration was determined by BCA assay. The solution corresponding to 200 μg proteins was then diluted to 4 M urea with 50 mM TEAB and digested in LysC at the enzyme:protein mass ratio of 1:100 for 4 hours at 37°C at 600 rpm. The mixtures were then diluted to 1 M urea with 50 mM TEAB and subjected to trypsin digestion (enzyme:protein mass ratio = 1:100) overnight at 37°C at 600 rpm.

### Sample preparation for total proteomics measurements

Worm lysates from cryomilling were resuspended in an equal volume of Buffer 5 (Buffer 1 + 4% SDS, 4 mM DTT, 8 mM EDTA, 2% (v/v) IGEPAL CA-630, 1:500 EDTA-free protease inhibitor tablet (Roche)), followed by incubation at 95°C for 20 min. The protein concentration was determined by BCA assay. 200 μg denatured protein samples were subjected to reduction, alkylation, protein purification, LysC and trypsin digestion as described above for the supernatant and pellet samples.

### Sample preparation and TMT labeling for mass spectrometry

Resultant peptides derived from all protocols listed above were desalted with a solidphase extraction (SPE) cartridge (Oasis HLB 1 cc Vac Cartridge, 10 mg sorbent, Waters Corp., USA), followed by lyophilization by freeze drying (Virtis, SP Scientific) overnight. The dried peptides were resuspended in 200 mM EPPS pH 8.5 and quantified by bicinchoninic acid assay using the quantitative colorimetric peptide assay from Thermo Fisher Scientific. 25 μg of peptides were subject to TMT labeling. For that, 28% of acetonitrile was added to each of the samples and labeled for 1 hour at room temperature with 60 μg of the corresponding TMTpro 16- or 18-plex isobaric labelling reagents (Thermo Fisher Scientific). Labeling scheme and information of the samples are included in Table S1. Reaction was quenched with 0.5% hydroxylamine for 15 min at room temperature. After labeling, 1.5 μL of each sample were pooled, desalted and analyzed on a Q-Exactive mass spectrometer (Thermo Fisher Scientific) to check labeling efficiency and corresponding mixing ratios. Final mixing volumes used for each of the samples to create the final pool were calculated based on the normalization factors listed in Table S1. Each pooled sample was desalted using a 50 mg Sep-Pak cartridge (Waters). Samples were dried, resuspended in 5% acetonitrile and 10 mM ammonium bicarbonate, pH 8.0 and subjected to basic pH reversed phase HPLC (bRPLC). All 96 fractions collected were combined into 24, 12 of which (not adjacent) were finally desalted before LC-MS/MS. For more information about the different TMT kits used and the labeling schemes used, see Table S1.

### Mass spectrometry analysis

All TMT included in this manuscript were analyzed by liquid chromatography-nano electrospray ionization-tandem mass spectrometry (LC-nESI-MS/MS) in an Orbitrap Fusion Lumos equipped with a high-Field Asymmetric Ion Mobility Spectrometry (FAIMS Pro) interface^69,70^ and coupled to a Proxeon NanoLC 1200 (Thermo Fisher Scientific). A 100 μm capillary column was packed with 35 cm of Accucore 150 resin (2.6 μm, 150 Å; Thermo Fisher Scientific). Peptides were separated at 525 nL/min flow rate using two buffers as mobile phases (Buffer A: 5% acetonitrile, 0.1% formic acid and Buffer B: 95% acetonitrile, 0.1% formic acid). Gradient lengths of each TMT analysis are included in Table S1. All MS parameters used for high resolution-MS2 (HR-MS2) quantifications for each TMT analysis are included in Table S1 (FAIMS compensation voltage -CV-, MS1 orbitrap resolution, scan range, MS1 maximum injection time, automatic gain control -AGC-, MS2 isolation window, higher-energy collision dissociation -HCD-, Orbitrap MS2 resolution, and MS2 maximum injection time and AGC).

### Proteomic database search

MS raw files were converted to mzXML using Monocle^71^ and processed using a Comet-based searching software pipeline^72,73^. Database searching included *C. Elegans* proteome (UP000001940) downloaded from Uniprot in October 2020. Most common contaminants and E. Coli proteome (UP000000558, downloaded from Uniprot in October 2020) were used as contaminants. Reversed sequences of all proteins were included in the search database for target-decoy FDR analysis^74,75^. Comet searches were performed using the following parameters: two missed cleavages and two variable modifications were allowed per peptide, precursor mass tolerance of 20 ppm, fragment ion mass tolerance of 0.02 Da, static modifications of cysteines (carboxyamidomethylation, +57.0215 Da), TMTpro labelling of lysines and N-termini of peptides (+304.2071 Da), and variable oxidation of methionine (+15.9949 Da). For LiP analysis (PK), semi-digested option was selected to allow one non-tryptic terminus in the peptides analyzed (half-tryptic). For all other analyses (TO, PEL and total), fully digested option was selected. Peptide-spectrum matches (PSM) filtering was performed using a linear discriminant analysis and adjusted to 1% FDR^74–76^. Proteins were assembled by principles of parsimony to produce the smallest set of proteins necessary to account for all observed peptides of each of the 3 experiments included in this manuscript (note that experiment one is composed by 3 TMT while experiments 2 and 3 are composed by 4 TMT each, Table S1). TMT reporter ion intensities were measured using a 0.003-Da window around the average m/z for each reporter ion (included in Table S1) in the MS2 scan. PSM with poor quality MS2 spectra were excluded from quantitation (<100 summed signal-to-noise -S/N- across all TMT channels and <0.7 precursor isolation specificity). Reporter ion intensities were adjusted to correct for the isotopic impurities of the different TMTpro reagents according to the manufacturer’s specifications (information of all TMT kits used is listed in Table S1). S/N measurements of peptides assigned to each protein were summed, and these values were normalized assuming equal protein loading for al samples in each TMT. Finally, each protein abundance was scaled to a percent of the total, such that the summed S/N for that protein across all channels equalled 100, thereby generating a relative abundance measurement. Number of total peptides and proteins identified and quantified are included in Table S1.

### Data analysis

For peptide PK accessibility analysis, fully- and half-tryptic peptides were used for quantitation. The total peptide intensities were normalized to be equal for each TMT channel (normalization factors in Table S1). To control for protein abundance changes between conditions, the peptide ratios were corrected by their corresponding protein ratios generated from the TO pipeline if significant (FDR < 0.05). For non-significant protein changes in the TO fraction, the correction factor is 1. The FDR was obtained from linear models for microarray (LIMMA) analysis^77^ using Benjamin-Hochberg correction for *P* values using R packages. The threshold for significant PK accessibility changes of peptides is |log_2_(fold change) | > 2 and FDR < 0.05.

For protein solubility (TO and aggregate) analysis, only fully-tryptic peptides were searched and used for quantitation. The total peptide intensities were normalized to be equal for each TMT channel (normalization factors in Table S1). To control for total protein abundance changes between conditions, the protein ratios were corrected by their corresponding protein ratios generated from the total protein abundance pipeline if significant (FDR < 0.05). For nonsignificant protein changes in the total fraction, the correction factor is 1. The FDR was obtained from LIMMA analysis using Benjamin-Hochberg correction for *P* values. The threshold for significant protein solubility changes is |log_2_(fold change) | > 2 and FDR < 0.05.

For total proteome abundance analysis, the total peptide intensities were normalized to be equal in each TMT channel. The threshold for significant protein total abundance changes is |log_2_(fold change) | > 2 and FDR < 0.05. The FDR was obtained from LIMMA analysis using Benjamin-Hochberg correction for *P* values.

### Bioinformatic analysis

For GO enrichment analysis for biological process, cellular component and molecular function, data were analyzed using GOATOOLS^78^. Fisher’s exact tests were performed with *P* values corrected using Benjamini-Hochberg multiple test correction. Results were filtered for GO terms with a fold enrichment over 2 unless stated otherwise and FDR less than 0.05. Protein interaction networks were generated using Cytoscape 3.9.1^79^ with the built-in String (v11.5) tool^80^ with a minimum required interaction score of 0.4 (medium confidence). Chaperone annotation was performed using the *C. elegans* chaperome list previously reported^43^. Clustal Omega^81,82^ was used to align the sequences of ribosomal subunits between *C. elegans* and human.

### Real-time quantitative polymerase chain reaction (RT-qPCR)

RNA was extracted and purified using Trizol and QIAGEN RNeasy Plus Micro kit. cDNA synthesis was performed using reverse transcriptase (BioRad) and RT-qPCR was performed using SYBR green PCR reaction mixture (BioRad) in a CFX384 Touch Real-Time PCR system (BioRad) with the following conditions: (15 s at 95°C, 20 s at 56°C, 30 s at 72°C) × 39 cycles. Relative expression was calculated from Cycle threshold values using the standard curve method. The relative mRNA levels of genes of interest were normalized to the housekeeping genes *cdc-42* and *rpb-2*.

### Statistical analysis

Statistical parameters are reported in the figures and corresponding figure legends. Figures were generated using the Seaborn package from Python v3.0. *P* values were considered significant at values less than 0.05 unless otherwise stated.

## Data availability

The mass spectrometry proteomics data have been deposited in the ProteomeXchange Consortium database via the PRIDE^83^ partner repository program with the dataset identifier PXDXXXX.

## SUPPORTING INFORMATION

This article contains supporting information (supplementary figures 1 – 6 and tables 1 – 8).

## ACKNOWLEDGEMENTS

We thank members of the Morimoto laboratory and Finley laboratory and the Program Project Grant for their scientific advice and comments on this work. X.S. was supported by postdoctoral awards from the National Ataxia Foundation and the BrightFocus Foundation for Alzheimer’s Disease Research (A2022023F). M.A.P. was supported by the Goldberg Fund Fellowship and the Miguel Servet Program (CP21/00017). Funding for this project was provided by P01 AG05447 from the National Institutes on Aging to R.I.M., D.F. and S.P.G.

## AUTHOR CONTRIBUTIONS

X.S. and R.I.M. conceived the project. X.S. and M.A.P. designed and performed the proteomics experiments. J.A.P. obtained mass spectra. X.S., M.A.P and R.I.M. designed the data analysis. X.S. and M.A.P. made figures with inputs from D.F. and R.I.M. X.S. and R.I.M wrote the original draft with edits from M.A.P. All authors revised and approved the manuscript.

