## Supplementary figures and images for "Global proteome metastability response in isogenic animals to missense mutations and polyglutamine expansions in aging"

### Supplementary Figure 1

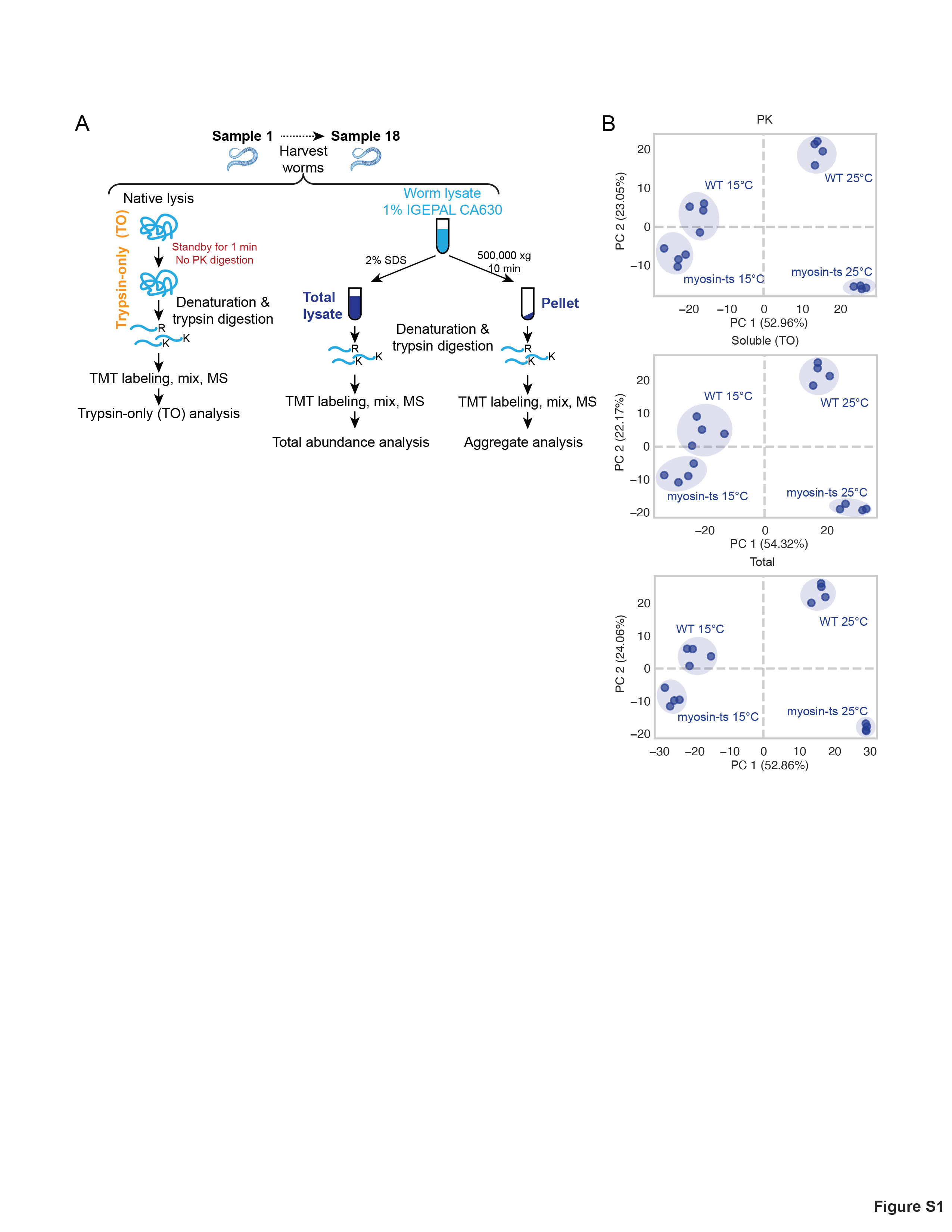

### Supplementary Figure 2

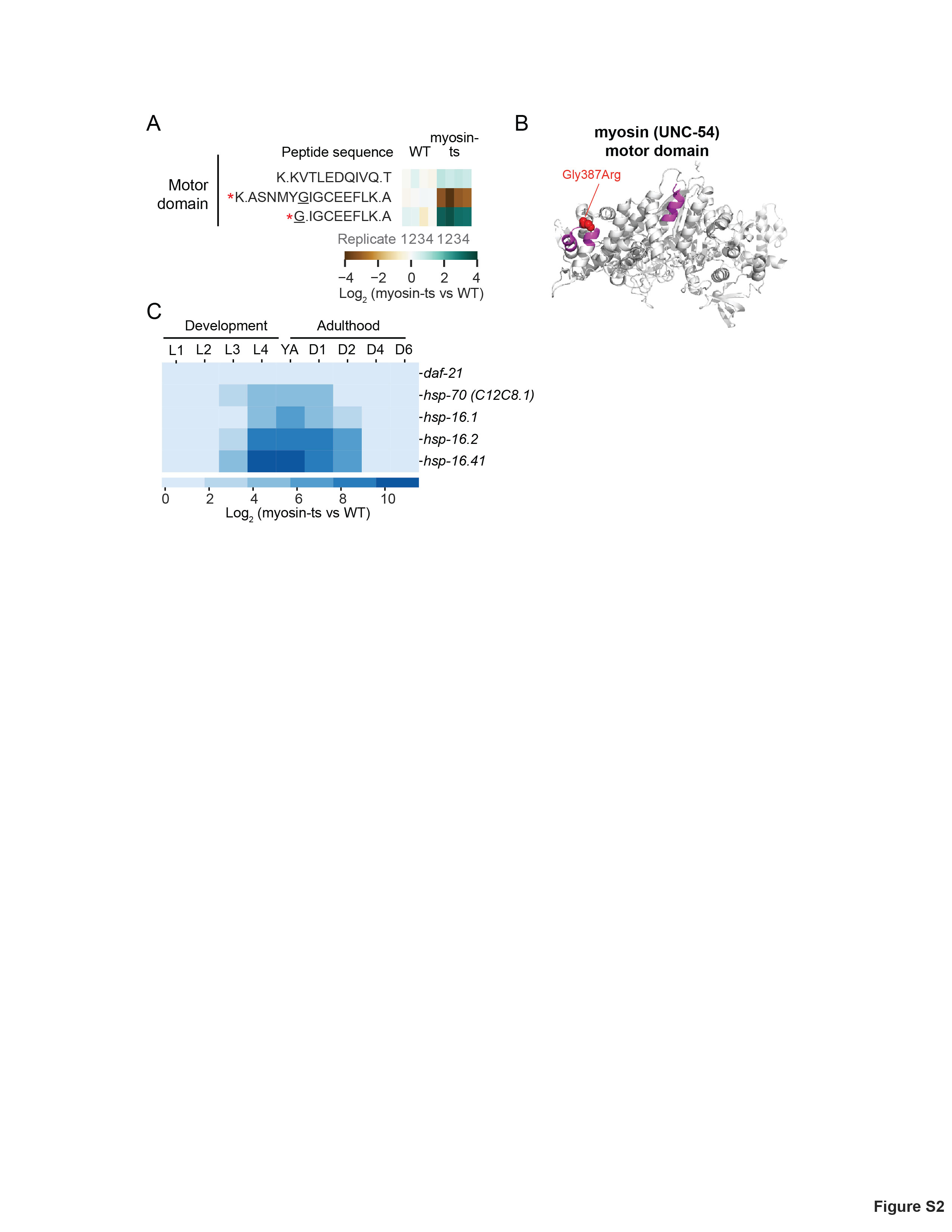

### Supplementary Figure 3

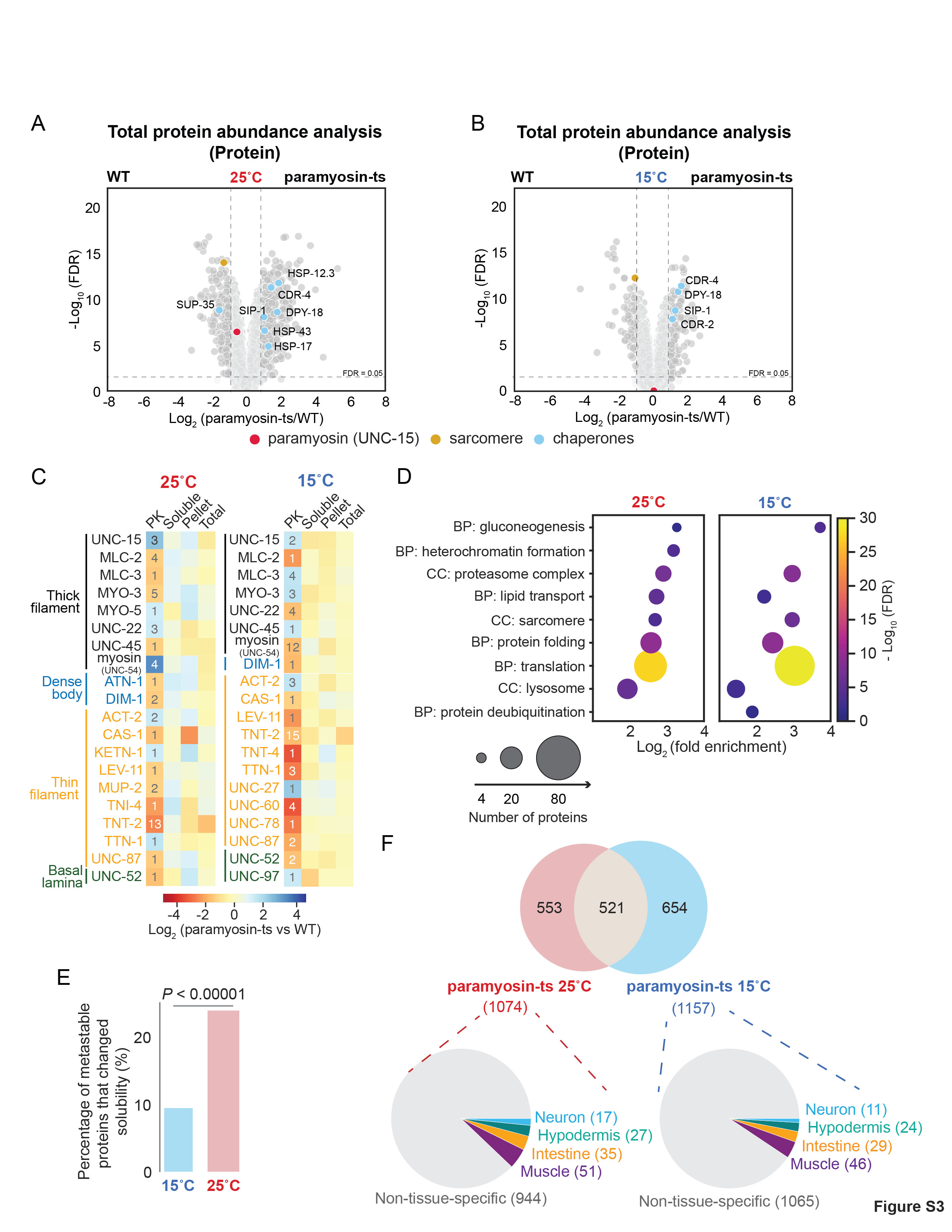

### Supplementary Figure 4

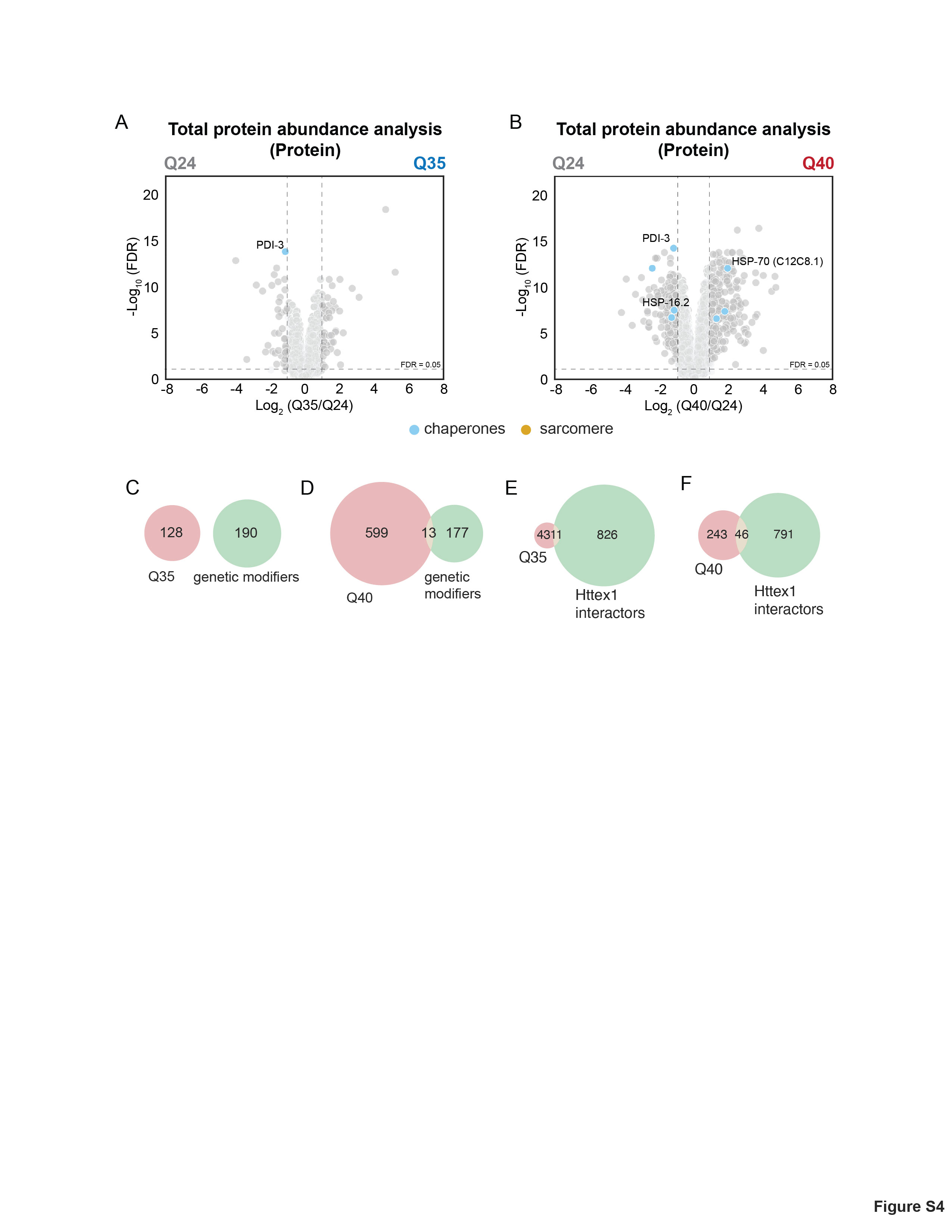

### Supplementary Figure 5

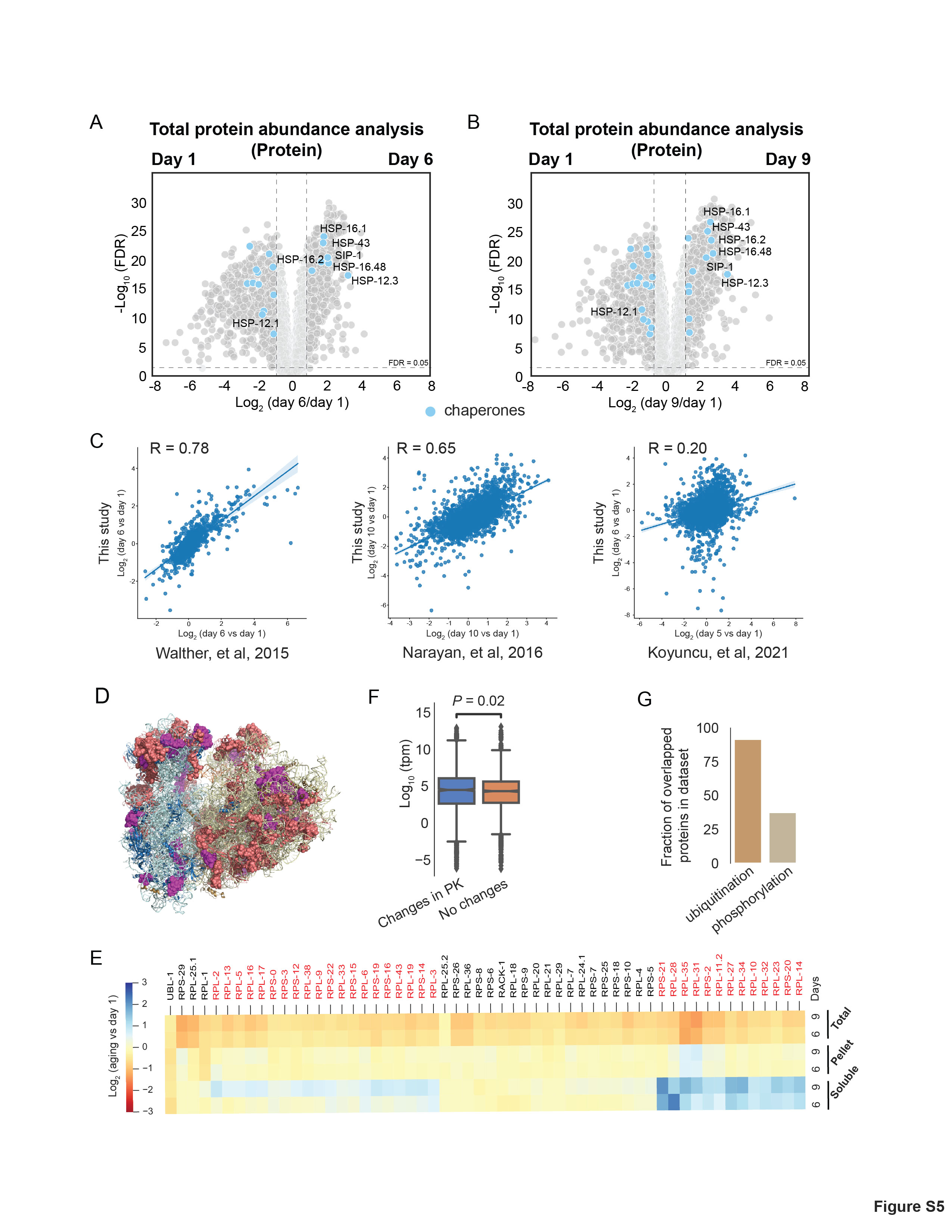

### Supplementary Figure 6

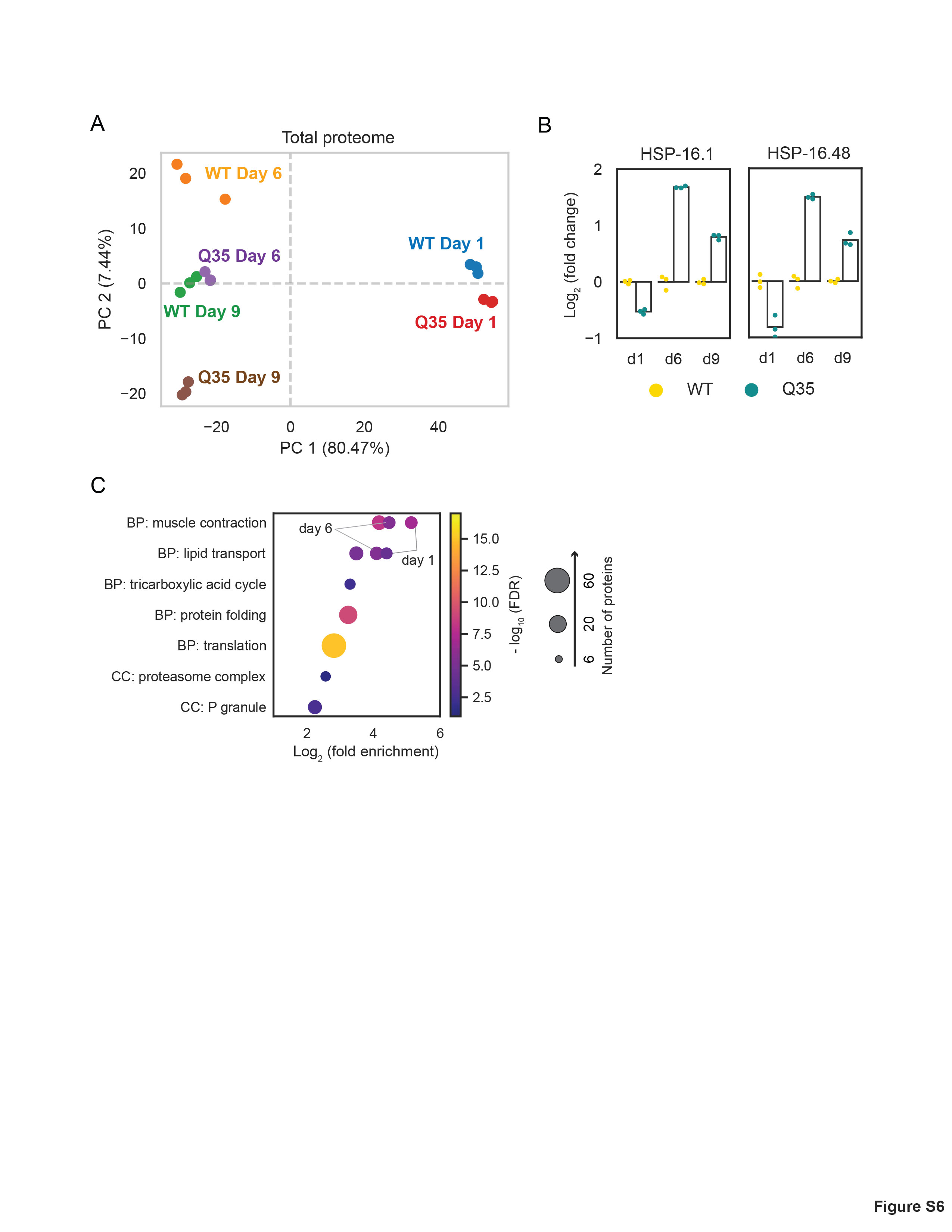
