## Supplementary information for "Global proteome metastability response in isogenic animals to missense mutations and polyglutamine expansions in aging"

**This file includes:**

Supplementary figure legends

Supplementary references

**Other supplementary materials for this manuscript include the following:**

Supplementary Table 1. MS and TMT information

Supplementary Table 2. Gene Ontology terms enriched among proteins changing PK accessibility in myosin-ts datasets

Supplementary Table 3. Tissue-specific mapping of proteins changing PK accessibility in ts datasets

Supplementary Table 4. Gene Ontology terms enriched among proteins changing PK accessibility in paramyosin-ts datasets

Supplementary Table 5. Gene Ontology terms enriched among proteins changing PK accessibility in polyQ datasets

Supplementary Table 6. Comparison of metastable proteins in polyQ strains with literature

Supplementary Table 7. Gene Ontology terms enriched among proteins changing PK accessibility in aging datasets

Supplementary Table 8. List of common proteins in seven conditions

**SUPPLEMENTARY FIGURE LEGENDS**

**Figure S1. Impact of a single ts missense mutation in myosin on the proteome conformational metastability.**

(A) Proteomic workflow for TMT-based quantitative measurements of changes in solubility (TO, left panel and aggregate, right panel) and total abundance (middle panel) across different conditions (up to 18 samples). To control for protein abundance changes between different conditions for PK accessibility analysis, same native protein lysates as Fig. 1B stood by for 1 min without PK digestion, followed by denaturation, trypsin digestion, TMT labelling, basic pH RP HPLC fractionation and MS analysis, called trypsin-only (TO) analysis. Protein ratios were calculated based on the sum of TMT reporter ion intensities of its associated peptides. As the starting material are from the soluble fraction, the TO analysis also reports protein abundance changes in the soluble fraction. As a complementary approach for solubility measurements, worm lysates were solubilized by 1% IGEPAL CA630, followed by ultracentrifugation at 500,000 g for 10 min at 4˚C to yield the detergent-insoluble pellet fraction. The pellets were then solubilized in 2% SDS, precipitated, resolubilized in 8 M Urea, trypsin digested, TMT labelled and subject to MS analysis, called aggregate analysis. As protein abundance changes in the soluble and pellet fraction are also affected by total protein abundance changes, to control for these, we also measured the total protein abundance changes between conditions using TMT proteomics, called total abundance analysis. From here on, protein ratios from soluble (TO) and pellet fractions represent those corrected by the ratios calculated from total abundance analysis. For correction details, see Materials and Methods session.

(B) Reproducibility of TMT-based proteomic analyses. Four biological replicates of myosin-ts and WT animals grown at the restrictive and permissive conditions at day 1 of adulthood were collected and analyzed for their changes in PK accessibility, solubility and total abundance. Principal component analysis (PCA) shows that different replicates of the same group cluster closely together in the two-dimensional plot (PC1 vs PC2), demonstrating the reproducibility among biological replicates.

**Figure S2. Impact of a single ts missense mutation on myosin conformation at the permissive condition.**

(A) Heatmap showing the myosin peptides with significantly differential PK accessibility between myosin-ts and WT animals from four biological replicates at the permissive condition. * represents the peptides that harbors the ts mutation (G387R, underlined). Note: the mutation site is cut by trypsin, therefore interfering the PK accessibility analysis.

(B) Crystal structure of the myosin motor domain (PDB: 6QDJ^1^) with all peptides with significant changes in PK accessibility colored in purple. The mutation site (G387R) is highlighted in red.

(C) Heatmap showing the mRNA abundance differences of chaperones (i.e., *daf-21*, *hsp-70 (C12C8.1)*, *hsp-16.1*, *hsp-16.2* and *hsp-16.41*) between myosin-ts and WT animals at various developmental stages (L1, L2, L3 and L4), young adult (YA), days 1, 2, 4 and 6 days of adulthood grown at the permissive condition.

**Figure S3. Impact of paramyosin-ts on the proteome at the restrictive and permissive conditions.**

(A – B) Volcano plots showing the total protein abundance changes in paramyosin-ts grown at the restrictive (A) and permissive (B) condition at day 1 of adulthood, compared with age-matched WT animals. Proteins with at least two-fold change and FDR < 0.05 (both thresholds indicated with dash lines) were considered significantly changed (colored in dark grey). The FDR was obtained from LIMMA analysis using Benjamin-Hochberg correction. Proteins were selectively annotated with their names. Among the significantly changed proteins, paramyosin (UNC-15), other sarcomere and chaperones were coloured in red, orange, and light blue, respectively.

(C) Dot plot showing the GO enrichment of BP and CC for proteins with significant changes in PK accessibility between paramyosin-ts and WT animals grown at the restrictive (left) and permissive (right) condition, respectively. Color indicates FDR values. Dot size indicates the number of proteins from the selected dataset that are included in the GO term. For full list of tissue-specific proteins, see Table S4.

(D) Heatmap showing the changes in PK accessibility, solubility (measured in the soluble (TO) and pellet fractions), and total abundance of sarcomeric proteins that showed significant changes in PK accessibility between paramyosin-ts and WT at the restrictive (left) and permissive (right) condition. The number shown under the PK category indicates the number of peptides with significant changes in PK accessibility. The median values of fold-changes of these peptides were used to represent the changes at the protein level. Protein names were colored based on their localisation on the sarcomere (thick filament: black, thin filament: orange, M-line: light blue, dense body: blue, and basal lamina: green).

(E) Bar plot showing the percentage of metastable proteins that changed their solubility at the permissive and restrictive conditions. Fisher’s exact test was performed to determine *P* value (annotated).

(F) Venn diagram showing the overlaps of proteins with significant changes in PK accessibility in paramyosin-ts at the restrictive and permissive conditions. The pie charts show the tissue-specific mapping of the metastable proteins. For the list of tissue-specific proteins, see Table S3.

**Figure S4. Total proteome changes in polyQ animals.**

(A – B) Volcano plots showing the total protein abundance changes in Q35 vs Q24 (A) and Q40 vs Q24 (B) animals. Proteins with at least two-fold change and FDR < 0.05 (both thresholds indicated with dash lines) were considered significantly changed (colored in dark grey). The FDR was obtained from LIMMA analysis using Benjamin-Hochberg correction. Proteins were selectively annotated with their names. Among the significantly changed proteins, chaperones were coloured in light blue.

(C – D) Overlap of proteins with significant changes in PK accessibility in Q35 (C) and Q40 (D) animals with genetic modifiers for polyQ aggregation.

(E – F) Overlap of proteins with significant changes in PK accessibility in Q35 (E) and Q40 (F) animals with interactors of polyQ-expanded huntingtin exon1 (Httex1) in Neuro2A cells. For the list of overlapped proteins, see Table S6.

**Figure S5. Total proteome changes in early aging in WT animals.**

(A – B) Volcano plots showing the total protein abundance changes in day 6 vs day 1 (A) and day 9 vs day 1 (B) WT animals. Proteins with at least two-fold change and FDR < 0.05 (both thresholds indicated with dash lines) were considered significantly changed (colored in dark grey). The FDR was obtained from LIMMA analysis using Benjamin-Hochberg correction. Proteins were selectively annotated with their names. Among the significantly changed proteins, chaperones were coloured in lightskyblue.

(C) Scatter plots of the protein fold changes in this study with published data (Walther et al. 2015; Narayan, et al, 2016; Koyuncu, et al, 2021; left to right), showing reproducibility of proteome changes between the studies. A strong correlation (R = 0.78 Pearson) was observed for comparison with Walther et al, 2015, suggesting the high reproducibility between these two studies for day 6 to day 1 comparison. A modest correlation (R = 0.65 Pearson) was observed for comparison with Narayan, et al, 2016. This is likely due to the use of 5'-fluorodeoxyuridine (FuDR) to avoid progeny contamination in their study and different isobaric labelling strategies (TMT vs SILAC). A low correlation (R = 0.20 Pearson) was observed for comparison with Koyuncu, et al, 2021. This is likely due to the use of FuDR in their study, different quantitation methods (TMT isobaric labelling vs label-free), as well as different timepoints examined (day 6 in this study vs day 5 in their study).

(D) Age-dependent changes in PK accessibility and solubility of ribosomal subunits. Cryo-EM structure of the human 80S ribosome (PDB: 6QZP^2^) with 40S subunits colored in blue, 18S rRNA in light blue, 60S subunits in wheat, 5S rRNA, 5.8 rRNA and 28S rRNA in light yellow, and tRNA in orange. All peptides with significant changes in day 9 vs day 1 in PK accessibility were colored in magenta, except for those with significant changes in solubility colored in red.

(E) Heatmap showing the solubility (soluble and pellet) and total abundance changes of ribosomal proteins that exhibited differential PK accessibility between days 1, 6 and 9 in WT animals. The subunits with changes in solubility were highlighted in red.

(F) Box plot comparing the translation levels (Stein, et al, 2022) of proteins with changes in PK accessibility and those without at day 6 of adulthood.

(G) Bar plot showing the fractions of proteins exhibiting age-dependent changes in ubiquitination (Koyuncu, et al, 2021) and phosphorylation (Wen-Jun, et al, 2021) among those with age-dependent changes in PK accessibility. Proteins compared were filtered for commonly identified proteins in the datasets.

**Figure S6. Acceleration of age-dependent proteome changes by Q35.**

(A) Three biological replicates of Q35 and WT animals grown to days 1, 6 and 9 of adulthood were collected and analyzed for their changes in total protein abundances. PCA showed that different replicates of the same group cluster closely together in the two-dimensional plot (PC1 vs PC2), demonstrating the reproducibility among biological replicates. This analysis also showed animals expressing Q35 at day 6 were clustered with WT animals at day 9.

(B) Bar plots showing total protein abundance changes of HSP-16.1 (left) and HSP-16.48 (right) between Q35 and WT animals at days 1, 6 and 9.

(C) Dot plot showing the GO enrichment of BP and CC for accumulated proteins with significant changes in PK accessibility between Q35 and WT at days 1, 6 and 9. GO terms enriched at day 1 and day 6 were annotated with the rest GO terms enriched at day 9. Dot size indicates the number of proteins included in the GO term and dot color indicates FDR values.
